# Resolving host and species boundaries for perithecia-producing nectriaceous fungi across the central Appalachian Mountains

**DOI:** 10.1101/2020.04.02.022061

**Authors:** Cameron M. Stauder, Nicole M. Utano, Matt T. Kasson

## Abstract

The Nectriaceae contains numerous canker pathogens. Due to scarcity of ascomata on many hosts, comprehensive surveys are lacking. Here we characterize the diversity of perithecia-producing nectriaceous fungi across the central Appalachians. Ten species from twelve hosts were recovered including a novel *Corinectria* sp. from *Picea rubens. Neonectria ditissima* and *N. faginata* were most abundant and associated with *Fagus grandifolia* with beech bark disease (BBD). *N. ditissima* was also recovered from additional cankered hardwoods, including previously unreported *Acer spicatum, Ilex mucronata*, and *Sorbus americana*. Cross-pathogenicity inoculations of *N. ditissima* confirmed susceptibility of *Acer* and *Betula* spp. *Neonectria magnoliae* was recovered from cankered *Liriodendron tulipifera* and *Magnolia fraseri* and pathogenicity on *L. tulipifera* was confirmed. *Fusarium babinda* was consistently recovered from beech with BBD, although its role remains unclear. This survey provides a contemporary snapshot of Nectriaceae diversity across the Appalachian Mountains. The following nomenclatural changes are proposed: ***Neonectria magnoliae*** comb. nov.

## 1. Introduction

Members of the Nectriaceae occupy diverse ecological niches from mycoparasites to phytopathogens, with numerous genera and species implicated in causing annual and perennial cankers on diverse woody plant hosts with varying degrees of host specificity. One such canker disease, beech bark disease (BBD), is a disease complex occurring across the range of American beech (*Fagus grandifolia* Ehrh.) in North America and European beech (*Fagus sylvatica* L.) in Europe. The disease requires prior infestation by an exotic scale insect (*Cryptococcus fagisuga* Lind.), which predisposes the host bark tissues to subsequent invasion by predominantly one of two canker fungi: *Neonectria ditissima* ([Tul. & C. Tul.] Samuels & Rossman) and either *N. faginata* ([Lohman, Watson, & Ayres] Castl. & Watson) on American beech or *N. coccinea* ([Pers.] Rossman and Samuels) (Houston, 1994b; Thomsen et al., 1949). In North America, BBD was first observed in Halifax, Nova Scotia, Canada around 1890 and presently continues to spread throughout the continuous range of American beech (Hewitt, 1914; Erhlich, 1934; Cale et al., 2017). Other members of Nectriaceae have occasionally been associated with BBD, including *Bionectria ochroleuca* ([Schwein.] Schroers & Samuels) and *Fusarium* spp., but their roles, if any, in BBD are not well understood (Cotter and Blanchard, 1982; Houston et al., 1987; Kasson and Livingston, 2009).

In addition to infecting scale-infested beech, *Neonectria ditissima* (formerly *N. galligena*) is well-known perennial target canker pathogen on many hardwood tree species often co-occurring in beech forests impacted by BBD (Lohman and Watson, 1943; Spaulding et al., 1936; Booth, 1967). Unlike BBD, which requires predisposition of host tissues by beech scale or possibly *Xylococculus betulae* (Cale et al., 2015), no insect partner has yet been established as a causal factor for *N. ditissima* infection on non-beech hosts. Based on previous pathogenicity studies, *N. ditissima* strains occurring in Eastern North America does not exhibit host specificity (Plante and Bernier, 1997), allowing unfettered interactions between inoculum produced from cankers on beech and non-beech hosts. Although this has not been confirmed among all dominant host species co-occurring with beech in BBD areas, namely black birch (*Betula lenta* L.) and striped maple (*Acer pensylvanicum* L.), mating barriers do not exist among these strains. This indicates there is likely no host specificity given an outcrossing population of strains occurring on varying host species (Stauder et al., 2020). Further investigations are warranted as unknown host specificity and mating barriers could theoretically influence the population dynamics of *N. ditissima* strains participating in the BBD complex by limiting or disrupting gene flow.

In contrast to *N. ditissima, N. faginata* (formerly *N. coccinea* var. *faginata*) has only been observed causing cankers following *C. fagisuga* infestation on American beech trees (Castlebury et al., 2006), leaving questions regarding its origin and ecological niche, if any, outside of BBD. Other closely related *Neonectria* species such as *N. punicea* also occur in eastern North America within northern hardwood forests and exhibit morphological attributes that overlap with *N. faginata* including ascospore size (Booth, 1959; Castlebury et al., 2006), which has historically served as a diagnostic measure in previous studies (Houston, 1994a; Kasson and Livingston, 2009; Lohman and Watson, 1943). As such, previous misidentifications of *Neonectria* spp. on beech and non-beech hosts are plausible when relying on morphological attributes, possibly underestimating diversity (for example see Fig 2 in Kasson and Livingston, 2009). Likewise, the close relationships among *Neonectria* spp., coupled with the lack of sequence data in repositories such as NCBI, has also presented species identification challenges, especially for rarer species, which further complicates identification. For example, our recent phylogenetic analyses of mating type gene sequences for *Nectria magnoliae* from *Liriodendron tulipifera* L. revealed this species formed a genealogically exclusive clade among other *Neonectria* species that was closely allied with *N. faginata* (Stauder et al., 2020). This supports previous findings by Gräfenhan and colleagues (2011), who showed but did not discuss that *N. ditissima* and *N. magnoliae* were genealogically exclusive. However, earlier work by Castlebury et al. (2006) concluded *N. magnoliae* was a synonym of *N. ditissima*. These findings emphasize the need for enhanced surveys to recover and phylogenetically resolve cryptic and understudied members of Nectriaceae as well as investigate the potential for cryptic reservoirs of known species including the possibility of *N. faginata* occurring on a non-beech host.

**Fig. 1:**
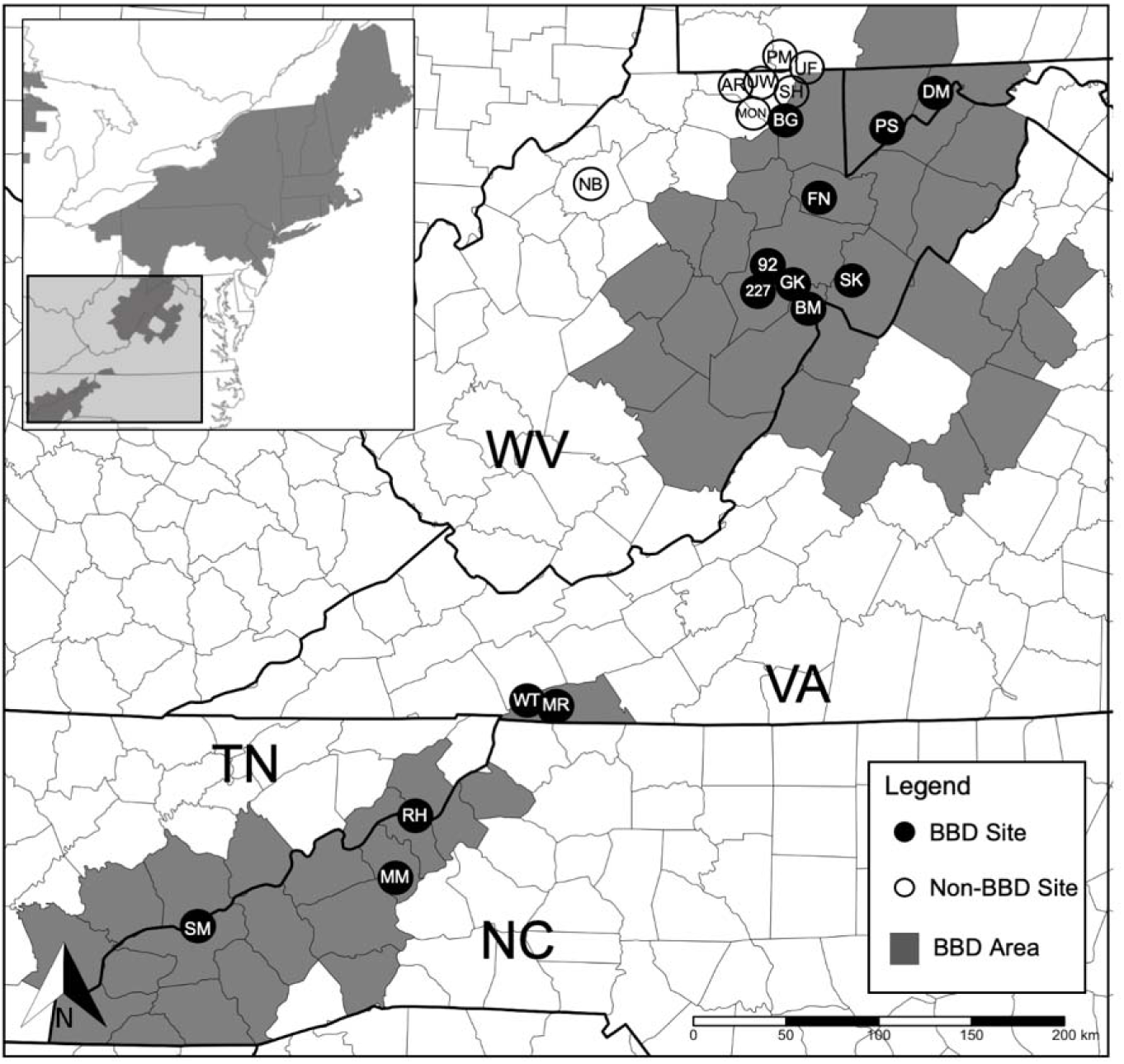
Sampling locations for members of Nectriaceae in the central Appalachian Mountains. Sites with confirmed beech bark disease (BBD) are designated by filled circles, sites lacking BBD are open circles, and counties where BBD has been previously reported are filled. BBD was first discovered in WV at site GK in 1981 (Mielke et al. 1982, Mielke and Houston, 1983). BBD distribution is based on a county level database (Blackburn, 2018).

**Fig. 2:**
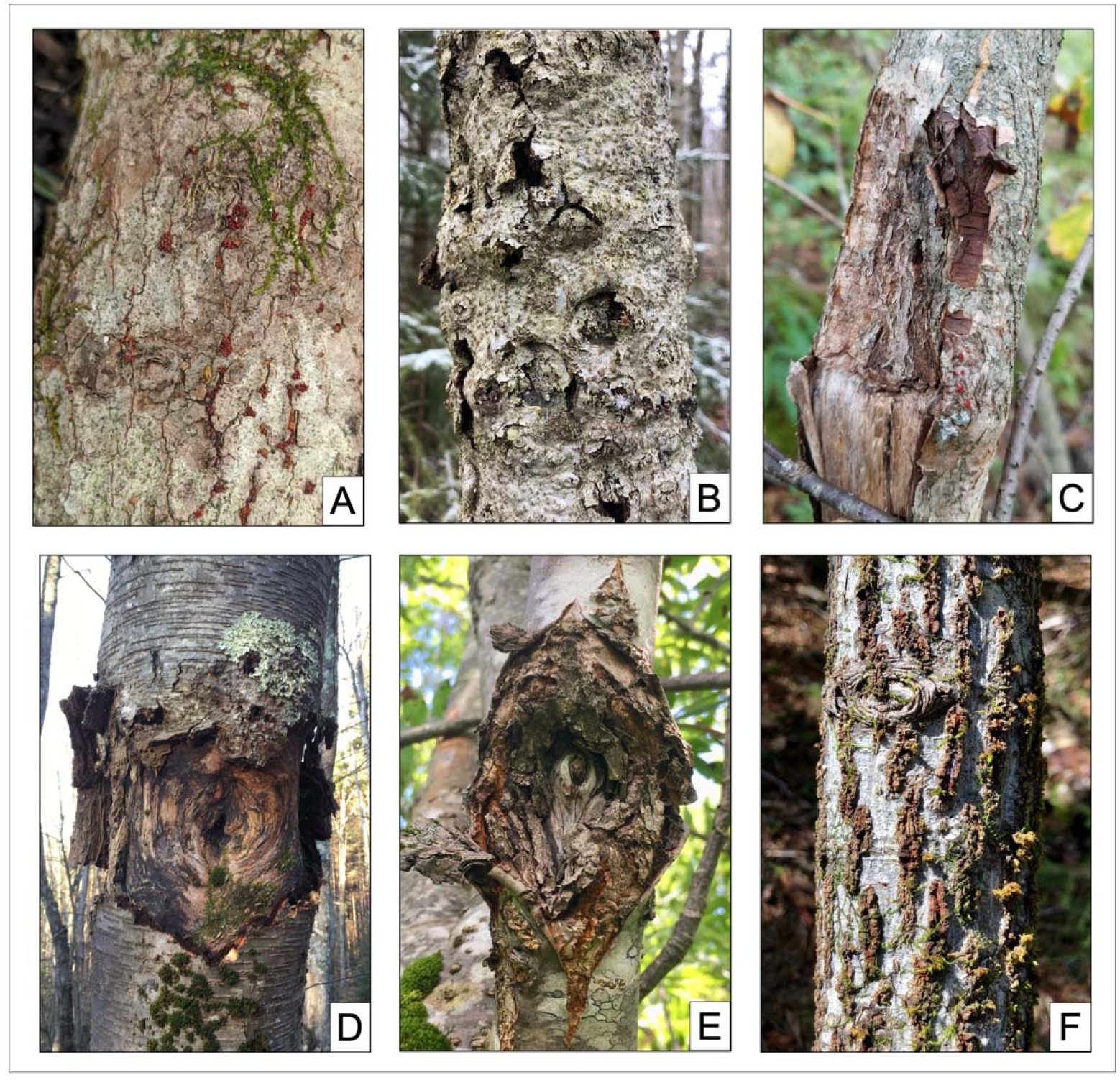
Example *Neonectria ditissima* canker photos for the following host species: A) mountain holly (*Ilex mucronate* at SK), B) American beech (*Fagus grandifolia* at GK), C) mountain maple (*Acer spicatum* at MM), D) black birch (*B. lenta* at UF), E) mountain ash (*Sorbus Americana* at WT), F) striped maple (*A. pensylvanicum* at SK)

This study sought to provide a greater understanding of the ecology and genetic relationships among *N. ditissima, N. faginata*, and allied fungi. The first objective of this study was to survey perithecia-producing members of Nectriaceae in American beech stands to identify possible native reservoirs of *N. faginata*. This is important given the uncertainty surrounding the origin of *N. faginata* and the potential for previous misidentifications of nectriaceous fungi recovered from non-beech hosts. The second objective was to resolve phylogenetic relationships among described and possibly undescribed members of Nectriaceae recovered from forests across the central Appalachian Mountains. This was critical given both the significant undersampling of nectriaceous fungi in these regions, the resulting the lack of sequence data, and/or contradictory evidence regarding certain known members of Nectriaceae, such as *Nectria magnoliae*. A third objective was to test host specificity of *N. ditissima, N. faginata, and N. magnoliae* strains isolated from American beech trees and tulip poplar. Together these aims sought to provide contemporary insights into the true diversity, phylogenetic relationships, and ecological niches of perithecia-producing nectriaceous fungi across the central Appalachian Mountains.

## 2. Materials and Methods

### 2.1. Site Selection

Sampling locations were selected based on previous reports of cankers and/or perithecia production on woody hosts, accessibility, and ability to secure sampling permits. In total, 13 beech bark disease (BBD) and eight non-BBD sampling locations were selected across WV, MD, VA, PA, TN and NC (Fig. 1; Supplemental Table 1). Four of the eight non-BBD sites served for sampling nectriaceous fungi from non-beech hosts, while the other four served as sites where beech tissue samples were collected from stands with no history of BBD. No defined sampling areas (*i.e*. plots) were established, but instead, trees were sampled opportunistically until sampling needs were met based on the sampling procedure described below. Sites GK, MM, and SM had five, two, and four geographically separated discrete sampling areas (Supplemental Table 1). All other sites are represented by a single sampling area.

Three WV sites (BM, GK, FR) were originally confirmed as having BBD at least 35 years prior to the initiation of this study (Mielke et al., 1982, Houston, 1994a). Sites designated BG, DM, and PSF represented more recent BBD outbreaks (∼ 15 years) as determined by various natural resource agency surveys in WV and MD. All other sites were determined to have likely become afflicted with BBD between 5 to 25 years prior (Morin et al., 2007). Four of seven sites lacking BBD (NB, PM, AR, UW) were selected for asymptomatic beech sampling based on a lack of previous reports, and all trees were visually confirmed to be absent of BBD-associated signs and symptoms prior to sampling.

### 2.2. Sampling Procedure

At each BBD site, American beech trees symptomatic for BBD and non-beech trees with perennial target cankers were identified and sampled. For approximately five trees of each species per site, a maximum of four bark disks harboring fresh perithecia were excised with a 1-cm diameter steel punch and stored in a microtiter plates. Additionally, bark tissue samples (∼1 mm diameter plugs; ∼8 samples/tree; up to 7 beech trees/site) were taken from symptomatic and asymptomatic trees at BBD and non-BBD sites, respectively. All samples were stored on ice until arriving at the laboratory where samples were stored at −20 °C until processing.

### 2.3. Sample Processing

To process each bark disk, up to five perithecia were removed with a sterile scalpel and cleaned by gently pushing each perithecium through sterile agar to remove debris. Each perithecium was squashed in a 1.5 ml microcentrifuge tube containing 1 ml of sterile H_2_O with a micropestle, vortexed for 15 seconds, and then 300 µl of the spore suspension was spread with a cell spreader on glucose-yeast extract agar plates amended with streptomycin sulfate (10 mg/1000 ml) and tetracycline hydrochloride (100 mg/1000 ml) antibiotics (GYE/A). Within 48 hours of plating, five germinating ascospores were subcultured to a new plate, and one isolate was selected for storage at approximately –20 °C on glass filter paper. All micro-sampled bark plugs were surface disinfested by soaking for 14 minutes in a 1:10 commercial bleach-water solution then up to four samples were placed onto each GYE/A agar plate. Resulting fungi of interest were subcultured individually to new plates then stored as previously described.

### 2.4. Species Identification

Recovered isolates were grouped and tentatively identified based on colony and macroconidia morphology and a subset of isolates were selected to be confirmed by DNA sequencing. Genomic DNA was extracted from isolates using a Wizard® kit (Promega, Madison, WI, USA) and suspended in 75 µl Tris-EDTA (TE) buffer (Amresco, Solon, OH, USA). PCR reactions were performed for the fungal barcoding genes internal transcribed spacer region (ITS), translation elongation factor 1-alpha (EF-1α), 28S rDNA (LSU), and β-tubulin (BTUB). PCR products were generated in 25 µl reactions containing 12.5 µl Bioline PCR Master Mix (Bioline USA Inc, Taunton, MA), 10.0 µl H_2_O, 1.5 µl purified DNA, and 1.0 µl each of forward and reverse primers (Integrated DNA Technologies, Coralville, IA, USA). All primers, protocols, and their sources are listed in Supplemental Table 2. Positive reactions identified via gel electrophoresis were prepared for sequencing using ExoSAP-IT (Affymetrix, Santa Clara, CA, USA) according to the manufacturer’s recommendations and Sanger sequenced using forward PCR primers (Eurofins, Huntsville, AL, USA). Resultant sequences were used to identify species using BLASTn searches, and the best match in the NCBI database was selected for the isolate’s identity.

### 2.5. Phylogenetic Analyses

For all sequences included in our phylogenetic analyses, chromatograms were assessed and clipped using CodonCode Aligner v. 5.1.5. Sequences were then manually corrected for nucleotide misreads by referencing their respective chromatograms. To examine phylogenetic relationships among collected members of Nectriaceae, single-gene and concatenated phylogenetic trees were constructed for all *Corinectria, Fusarium*, and *Neonectria* species observed in this survey along with additional reference sequences for selected members of Nectriaceae available from NCBI Genbank (Table 3). The other members of the Nectriaceae recovered either lacked sampling depth or adequate reference sequences for our loci of interest to permit meaningful phylogenetic analysis. As such, only BLASTn searches of individual loci were conducted for these species with representative sequences deposited into NCBI Genbank. For *Corinectria, Fusarium*, and *Neonectria* species, each gene was aligned using MAFFT (Katoh and Stanley, 2013) on the Guidance 2.0 server (Landan and Graur, 2008; Sela et al., 2015). Individual resides with Guidance scores <0.5 were masked (Macias et al., 2020). A concatenated sequence was generated from single gene alignments using the web tool FaBox (Villesen, 2007).

**Table 1:**
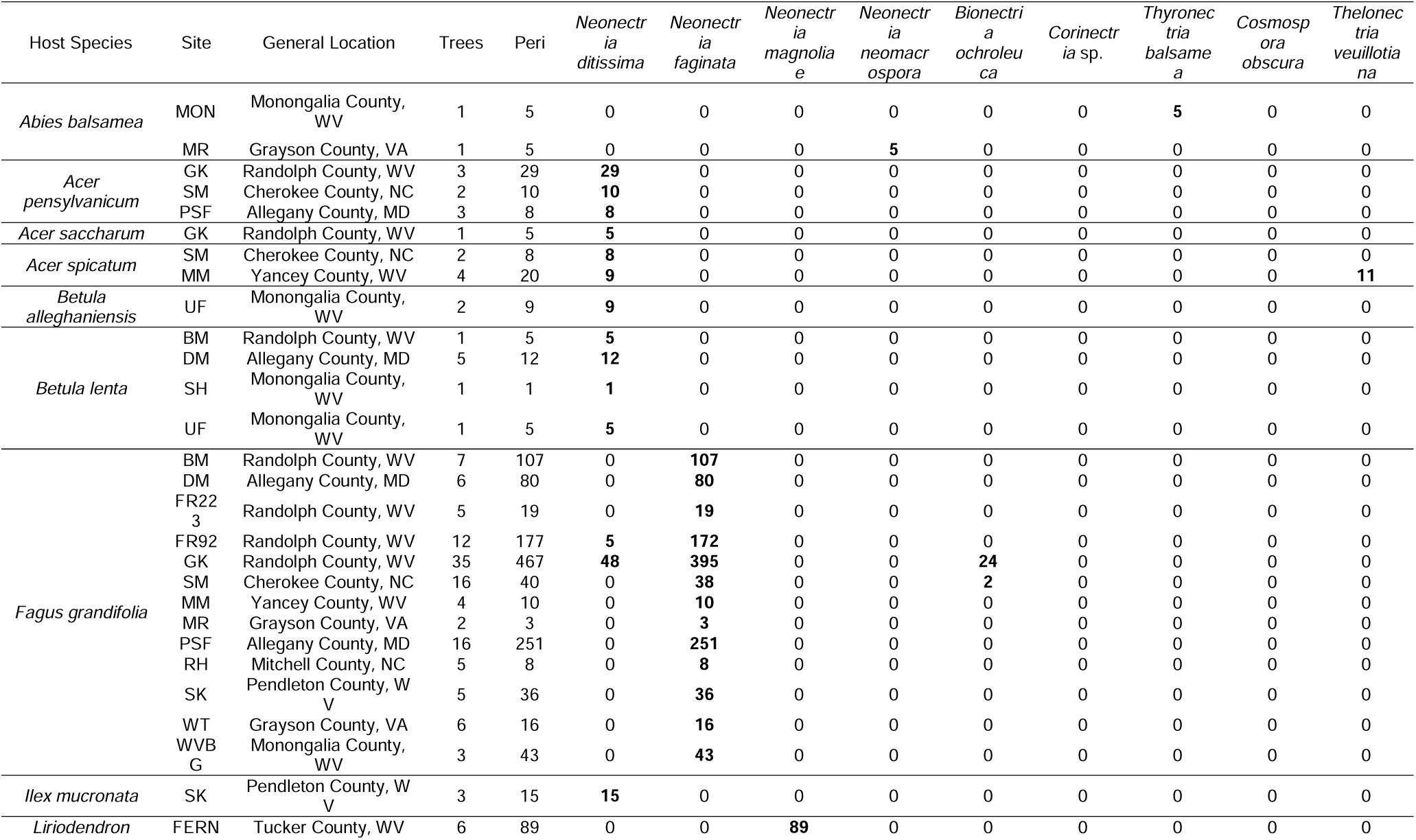
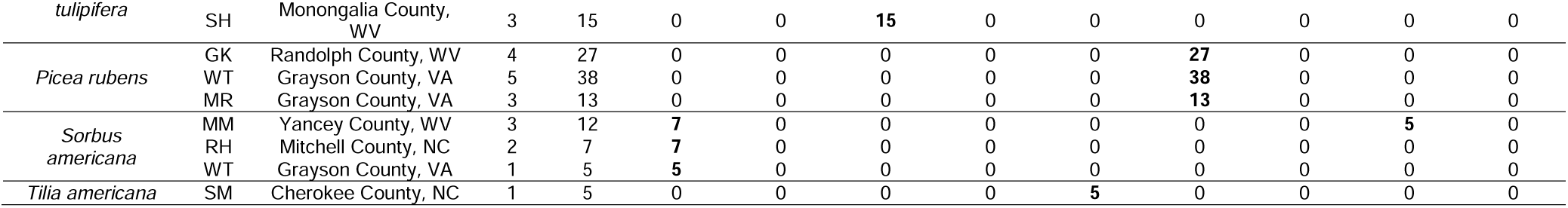
Survey sample collection summary by host tree species. Values under each fungal species represent the number of sampled perithecia yielding an isolate of that fungus. In total, 1,605 perithecia yielded nine fungal species from ten host tree species at 17 geographically separated sites.

**Table 2:** Isolate identification by BLASTn searches using ITS sequences derived from selected representatives of dominant morphologies. Percent identity values represent the lowest identity value among the sequences for each fungal species.

| Species | Isolate Count | Closest tblastn match | % identity |
| --- | --- | --- | --- |
| *Neonectria ditissima* | 17* | *Neonectria ditissima* | ≥ 98.29% |
| *Neonectria faginata* | 5 | *Neonectria faginata* | ≥ 99.02% |
| *Neonectria magnoliae* | 11 | *Nectria magnoliae* | ≥ 98.38% |
| *Neonectria neomacrospora* | 2 | *Neonectria neomacrospora* | 99.2% |
| *Bionectria ochroleuca* | 2 | *Bionectria ochroleuca* | ≥ 98.5% |
| *Corinectria* sp. | 5 | *Corinectria fuckeliana* | ≥ 94.8% |
| *Thyronectria balsamea* | 2 | *Thyronectria balsamea* | ≥ 99.44% |
| *Cosmospora obscura* | 2 | *Cosmospora obscura* | ≥ 99.04% |
| *Thelonectria veuillotiana* | 4 | *Thelonectria veuillotiana* | ≥ 99.64% |
| *Fusarium babinda* | 4 | *Fusarium babinda* | ≥ 99.8% |

\*Additional *N. ditissima* samples sequenced to represent isolates from varied host species

**Table 3:** Accession table for sequences used in the phylogenetic analyses of this study. Sequences were either acquired from NCBI or generated as part of this study. Host species and location are also provided where available.

| Species | Strain ID | Host | Locality | BT | LSU | EF1 | ITS |
| --- | --- | --- | --- | --- | --- | --- | --- |
| *Fusarium babinda* | FbFg001 | *Fagus grandifolia* | Allegany County, MD, USA | TBD | TBD | TBD | TBD |
| *F. babinda* | FbFg002 | *F. grandifolia* | Randolph County, WV, USA | TBD | TBD | TBD | TBD |
| *F. babinda* | FbFg003 | *F. grandifolia* | Allegany County, MD, USA | TBD | TBD | TBD | TBD |
| *F. babinda* | FbFg004 | *F. grandifolia* | Monongalia County, WV, USA | TBD | TBD | TBD | TBD |
| *F. concolor* | NRRL 13994 | *Hordeum* sp. | Uruguay | U61549.1 | U61653.1 | --- | U61679.1 |
| *F. concolor* | BCCM9748 | *Homo sapiens* | Spain | KJ125965.1 | KJ126557.1 | --- | KJ125669.1 |
| *Neonectria coccinea* | AF3707 | *F. sylvatica* | Slovakia | TBD | TBD | --- | TBD |
| *N. coccinea* | AR3691 | *F. sylvatica* | Romania | TBD | TBD | --- | TBD |
| *N. coccinea* | CBS 118916 | *F. sylvatica* | Romania | DQ789840.1 | KC660601.1 | KC660442.1 | KC660505.1 |
| *N. coccinea* | CBS 119158 | *Fagus* sp. | Germany | KC660727.1 | KC660620.1 | JF268734.1 | KC660521.1 |
| *N. confusa* | JL2009a-5741 | --- | --- | JF268721.1 | --- | JF268736.1 | JF268760.1 |
| *N. confusa* | CBS 127484 | --- | China | KM515886.1 | KM515933.1 | --- | KM515889.1 |
| *N. ditissima* | NdBa001 | *Betula alleghaniensis* | Monongalia County, WV, USA | TBD | TBD | TBD | TBD |
| *N. ditissima* | NdSam001 | *Sorbus americana* | Grayson County, VA, USA | TBD | TBD | TBD | TBD |
| *N. ditissima* | NdAp002 | *Acer pensylvanicum* | Swain County, NC, USA | TBD | TBD | TBD | TBD |
| *N. ditissima* | NdAs002 | *Acer spicatum* | Yancey County, WV, USA | TBD | TBD | TBD | TBD |
| *N. ditissima* | NdII001 | *Ilex mucronata* | Pendleton County, WV, USA | TBD | TBD | TBD | TBD |
| *N. ditissima* | NdBI002 | *Betula lenta* | Allegany County, MD, USA | TBD | TBD | TBD | TBD |
| *N. ditissima* | NdAsa001 | *Acer saccharum* | Randolph County, WV, USA | TBD | TBD | TBD | TBD |
| *N. ditissima* | NdFg002 | *F. grandifolia* | Randolph County, WV, USA | TBD | --- | TBD | TBD |
| *N. ditissima* | NdFg006 | *F. grandifolia* | Randolph County, WV, USA | TBD | TBD | TBD | TBD |
| *N. ditissima* | NdFg001 | *F. grandifolia* | Randolph County, WV, USA | TBD | TBD | TBD | TBD |
| *N. ditissima* | CBS 100.316 | --- | Ireland | HM352864.1 | HM364311.1 | HM364350.1 | HM364298.1 |
| *N. ditissima* | CBS 226.31 | *F. sylvatica* | Tharandt, Germany | DQ789869.1 | AY677330.1 | JF735783.1 | JF735309.1 |
| *N. faginata* | CBS 118938 | *F. grandifolia* | PA, USA | DQ789846.1 | KC660614.1 | KC660446.1 | KC660508.1 |
| *N. faginata* | CBS 217.67 | *F. grandifolia* | New Brunswick, Canada | JF268730.1 | HQ840382.1 | JF268746.1 | HQ840385.1 |
| *N. faginata* | NfFg014 | *F. grandifolia* | Allegany County, MD, USA | TBD | TBD | TBD | TBD |
| *N. faginata* | NfFg007 | *F. grandifolia* | Swain County, NC, USA | TBD | TBD | TBD | TBD |
| *N. faginata* | NfFg005 | *F. grandifolia* | Randolph County, WV, USA | TBD | TBD | --- | TBD |
| *N. faginata* | NfFg015 | *F. grandifolia* | Monongalia County, WV, USA | TBD | TBD | --- | TBD |
| *N. magnoliae* | NmLt001 | *Liriodendron tulipifera* | Tucker County, WV, USA | TBD | TBD | TBD | TBD |
| *N. magnoliae* | NmLt002 | *L. tulipifera* | Tucker County, WV, USA | TBD | --- | TBD | TBD |
| *N. magnoliae* | NmLt003 | *L. tulipifera* | Monongalia County, WV, USA | TBD | TBD | TBD | TBD |
| *N. magnoliae* | NmLt004 | *L. tulipifera* | Monongalia County, WV, USA | TBD | --- | TBD | TBD |
| *N. magnoliae* | NmMf001 | *Magnolia fraseri* | Randolph County, WV, USA | TBD | TBD | TBD | TBD |
| *N. punicea* | CBS 119532 | *F. sylvatica* | Slovakia | KC660728.1 | KC660563.1 | KC660439.1 | KC660503.1 |
| *N. punicea* | CBS 242.29 | *Rhamus* sp. | Germany | DQ789873.1 | MH866520.1 | DQ789730.1 | MH855054.1 |
| *N. neomacrospora* | NnAb002 | *Abies fraseri* | Grayson County, VA, USA | TBD | TBD | TBD | TBD |
| *N. neomacrospora* | NnAb001 | *A. fraseri* | Grayson County, VA, USA | TBD | TBD | TBD | TBD |
| *N. neomacrospora* | CBS 118985 | *Tsuga heterophylla* | Canada | DQ789890.1 | HQ840380.1 | JF268755.1 | HQ840389.1 |
| *N. neomacrospora* | CBS 118984 | *Abies balsamea* | Canada | DQ789882.1 | HQ840379.1 | JF735787.1 | HQ840388.1 |
| *N. hederæ* | IMI 058770a | *Hedera helix* | United Kingdom | DQ789895.1 | KC660617.1 | KC660461.1 | --- |
| *N. hederæ* | CBS 714.97 | *H. helix* | Netherlands | DQ789878.1 | KC660616.1 | KC660461.1 | QGQB00000000 |
| *N. ramulariae* | CBS 151.29 | *Malus sylvestris* | United Kingdom | DQ789863.1 | AY677333.1 | HM054091.1 | AY677291.1 |
| *N. ramulariae* | CBS 182.36 | *M. sylvestris* | --- | JF735439.1 | MH867276.1 | JF735792.1 | MH855762.1 |
| *Corinectria* sp. | CspPr004 | *Picea rubens* | Randolph County, WV,<br>USA | TBD | TBD | TBD | TBD |
| *Corinectria* sp. | CspPr007 | *P. rubens* | Randolph County, WV,<br>USA | TBD | TBD | TBD | TBD |
| *Corinectria* sp. | CspPr002 | *P. rubens* | Grayson County, VA,<br>USA | TBD | TBD | TBD | TBD |
| *Corinectria* sp. | CspPr008 | *P. rubens* | Grayson County, VA,<br>USA | --- | TBD | TBD | TBD |
| *C. tsugae* | CBS 788.69 | *T. heterophylla* | Canada | KM232020.1 | MH871201.1 | --- | KM231763.1 |
| *C. fuckeliana* | IMI 342668 | *Picea* sp. | Switzerland | KJ022340.1 | KJ022070.1 | KJ022404.1 | KJ022021.1 |
| *C. fuckeliana* | CBS 239.29 | *Picea sitchensis* | Scotland | DQ789871.1 | HQ840377.1 | JF268748.1 | HQ840386.1 |
| *C. constricta* | LASBE 266 | *Pinus radiata* | Chile | KY636417.1 | --- | KY636410.1 | --- |

For single gene and concatenated sequences, maximum-likelihood analyses were completed using MEGA v10.1.7 (Stecher et al., 2020), and Bayesian inference (BI) analyses were completed using MrBayes v. 3.2.7 (Ronquist et al., 2012). For ML analyses, the best-fit nucleotide substitution model was chosen using Model Test AICc scores in MEGA and 1000 bootstrap replicates were used. For BI analyses, MrBayes selected the best fit nucleotide selection model, but the rate of substitution was selected from the Model test AICc scores. The BI runs were stopped once the standard deviation of split frequencies fell below 0.01. These were then checked for convergence in Tracer v. 1.7.1 (Rambaut et al., 2018). Trees were prepared for publication using FigTree v. 1.4.4 (Rambaut, 2017) and Adobe Illustrator v. 24.1.

### 2.6. Morphological Measurements

Over the course of this survey, a species molecularly identified as *Nectria magnoliae* was recovered from tulip poplar (*Liriodendron tulipifera* L.) and Fraser magnolia (*Magnolia fraseri* Walter). Based on a recent study in which isolates from both of these hosts were included, this species was shown to form an independent clade with other members of *Neonectria* (Stauder et al., 2020). A previously published study also supported these relationships (Gräfenhan et al. 2011). Given these phylogenetic relationships, ascospore and conidia measurements were conducted to further characterize this species to permit comparisons with original descriptions provided by Lohman and Watson (1943).

For ascospore measurements, a total of 15 perithecia were processed. Three perithecia were sampled from each of five bark disks representing two geographically separated sites (FN = 3 disks; SH = 2 disks). A single perithecium was extracted from the bark disk using a sterile scalpel and squash-mounted on a slide with lactophenol plus cotton blue mountant. Length and width measurements were collected for 25 ascospores per perithecium. All measurements were taken with a Nikon Eclipse E600 compound microscope (Nikon Instruments, Melville, NY, USA) equipped with a Nikon Digital Sight DS-Ri1 microscope camera and Nikon NIS-Elements BR3.2 imaging software.

Conidia measurements were similarly conducted for both micro- and macro-conidia. Here, sporodochial masses were harvested from pure four-to six-week-old cultures using a sterile scalpel and mounted onto slides as described above. Both length and width measurements were taken for 50 microconidia were harvested from two isolates from each of three geographically separated locations (GK, SH, FN). Samples from SH and FN were collected from *L. tulipifera*, and samples from GK were collected from *M. fraseri*. Macroconidia were only found associated with one isolate collected from *M. fraseri* at GK and measured as described.

### 2.7. Pathogenicity Assays

Field inoculations were conducted to further investigate pathogenicity of *N. magnoliae* strains from *L. tulipifera* as well as *N. faginata* and *N. ditissima* strains from beech on birch, striped maple, and beech. For *N. magnoliae*, a single isolate (NmLt001) recovered from a natural infection on *L. tulipifera* was selected. One isolate of both *N. ditissima* (NdFg002) and *N. faginata* (NfFg005) recovered from American beech were also included for cross-pathogenicity testing. All study isolates were grown in pure culture on GYE for two-weeks at room temperature. Prior to inoculations, a sterile 1-cm steel punch was used to cut inoculation plugs along the growing edge of the colony. Negative control plugs were cut from sterile GYE plates.

Six each of tulip poplar (*L. tulipifera*), black birch (*Betula lenta*), yellow birch (*Betula alleghaniensis* Britt.), and striped maple (*Acer pensylvanicum*) trees, located on WVU University Forest were selected for inoculations. Additionally, five American beech trees were selected at this same location. Each tree received an inoculation with *N. ditissima, N. faginata*, and *N. magnoliae*. In addition to these fungal inoculations, a negative control inoculation with a sterile GYE agar plug was also performed on each study tree.

For each inoculations, a sterile 1-cm leather punch was used to excise bark tissue and create an inoculum reservoir. A colonized or sterile agar plug was then placed into the reservoir, and masking tape was applied over the wound to limit inoculum desiccation prior to infection. After six-months, bark tissue was excised from canker margins using a bone-marrow biopsy tool and placed in a 96-well microtiter dish. To ensure accurate canker measurements, a knife was used to remove bark tissue and reveal any underlying necrosis. Length and width measurements were then taken for each canker resulting from inoculation. Recovery of inoculant was achieved as previously described for bark tissue sample processing. All isolate identities were confirmed morphologically.

### 2.8. Statistical analyses

For pathogenicity measures, a one-way ANOVA was completed to check for differences in canker size and a Tukey-HSD post-hoc test was completed identify significant pairwise differences using the stats v3.6.2 package within R v 3.6.3 statistical software (R Core Team, 2020). All p-values <0.05 were considered significant.

## 3. Results

### 3.1. Survey of nectriaceous fungi

Perithecia of putative nectriaceous fungi were sampled from 180 trees across 17 sites in WV, MD, VA, NC, and TN (Table 1). Sampling included 12 tree species with black birch, red spruce, and mountain ash being the most abundantly sampled species besides America beech. A total of 1,605 sampled perithecia yielded nine fungal species spanning six genera. The majority of these samples were collected from American beech trees (n = 1,257 perithecia). The remaining 348 perithecia were sampled from one of the other 11 host species listed in Table 1. All isolates were grouped initially by morphology and a subset was identified with NCBI BLASTn searches using ITS barcoding sequences (Table 2; Supplemental Table 3). For NCBI blast hits, a >98% sequence identity was used as a cutoff threshold to confirm identification of previously described species.

Of samples collected from American beech trees, approximately 4.2% (53 out of 1,257) of sampled perithecia yielded isolates of *N. ditissima*, while 93.7% (1,178 out of 1,257) yielded *N. faginata*. The remaining 2.1% (26 out of 1,257) yielded *Bionectria ochroleuca*. Out of the 13 sites where American beech perithecia were sampled, *N. ditissima* was only recovered from locations FR92 and GK and represented 2.8% (5 out of 177) and 10.8% (48 out of 443) of the viable perithecia collected from those sites, respectively. In one instance, both *N. ditissima* and *N. faginata* were isolated from perithecia occurring on the same bark disk. *Bionectria ochroleuca* was recovered from 2 of 13 sites, GK and SM.

*N. ditissima* was also isolated from conspicuous cankers on a number of additional hardwood hosts including yellow birch (*Betula allegheniensis*), mountain maple (*Acer spicatum* Lam.) (Fig. 2C), black birch (*B. lenta*) (Fig. 2D), and mountain ash (*Sorbus americana* Marshall) (Fig. 2E) (Table 1). Additionally, *N. ditissima* was also recovered from the main stems of striped maple (*A. pensylvanicum*) and mountain holly (*Ilex mucronata* (L.) M.Powell, Savol., & S.Andrews), though the cankers were inconspicuous and mostly lacking callous ridges (Fig. 2A and 2F) (Table 1; Fig. 2; Fig. 3A). *Bionectria ochroleuca* was recovered from American beech and American basswood (*Tilia americanum* L.) (Fig. 3G). Three additional members of Nectriaceae were also recovered, albeit less frequently, during this survey on tree species found co-occurring with beech in BBD areas or nearby. These included *N. magnoliae* on tulip poplar (*L. tulipifera*) and Fraser magnolia (*Magnolia fraseri*) (Fig. 6), *Thelonectria veuillotiana* ([Roum. & Sacc.] P. Chaverri & Salgado) on mountain maple (*A. spitacum*) and mountain ash (*S. americana*) (Fig. 3F), and *Cosmospora obscura* (Rossman & Samuels) on mountain ash (Fig. 3H).

**Fig. 3:**
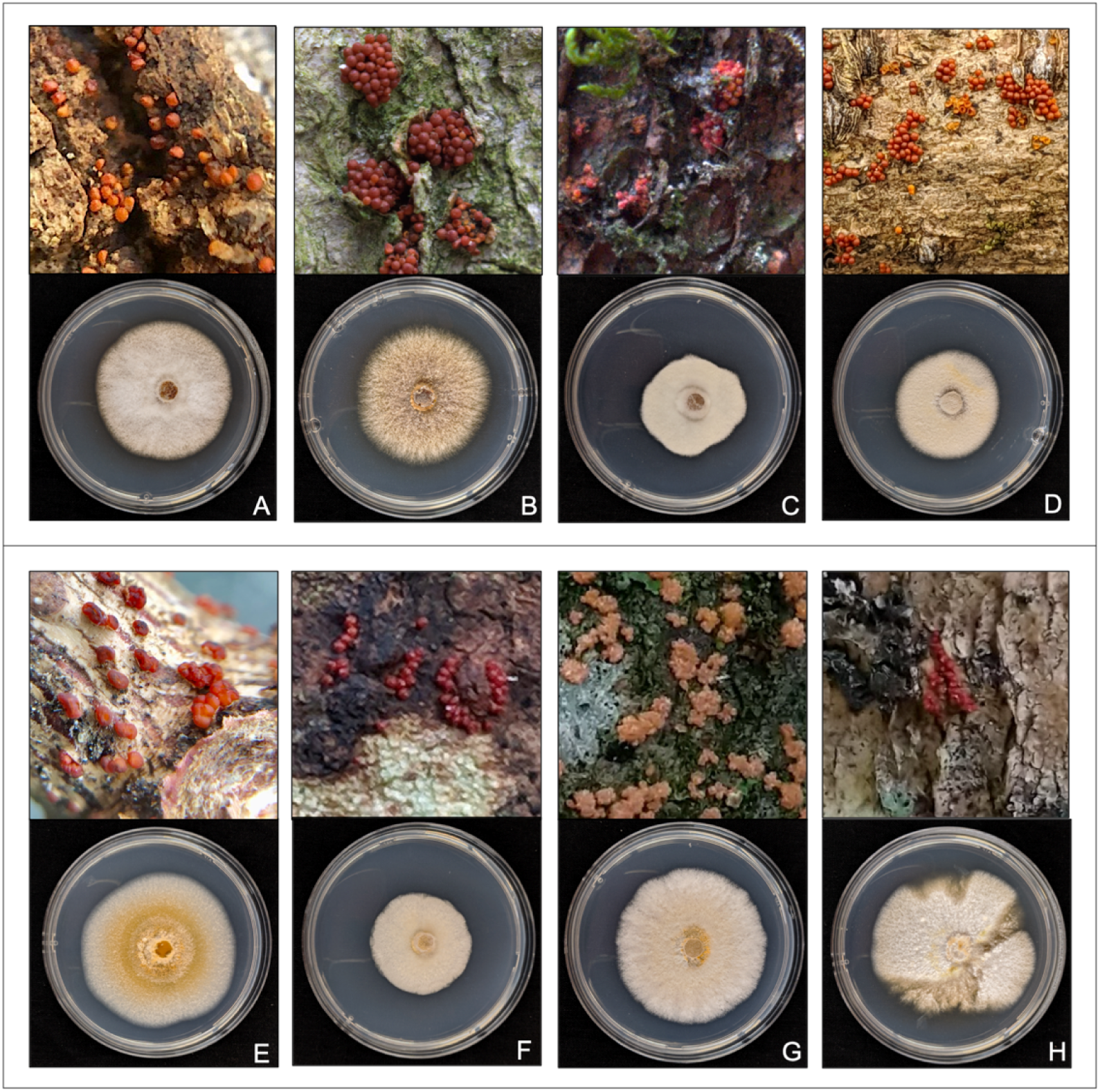
Diversity of Nectriaceae recovered across the central Appalachian Mountains on natural substrate and in culture: A) *Neonectria ditissima* (shown on striped maple at SK), B) *N. faginata* (on American beech at BG), C) *N. neomacrospora* (on Fraser fir at MR), D) *Corinectria* sp. (on red spruce at GK), E) *Thryonectria balsamea* (on balsam fir at MON), F) *Thelonectria veuillotiana* (on mountain maple at MM), G) *Bionectria ochroleuca* (on American beech at SM), and H) *Cosmospora obscura* (on mountain ash at MM).

**Fig. 4:**
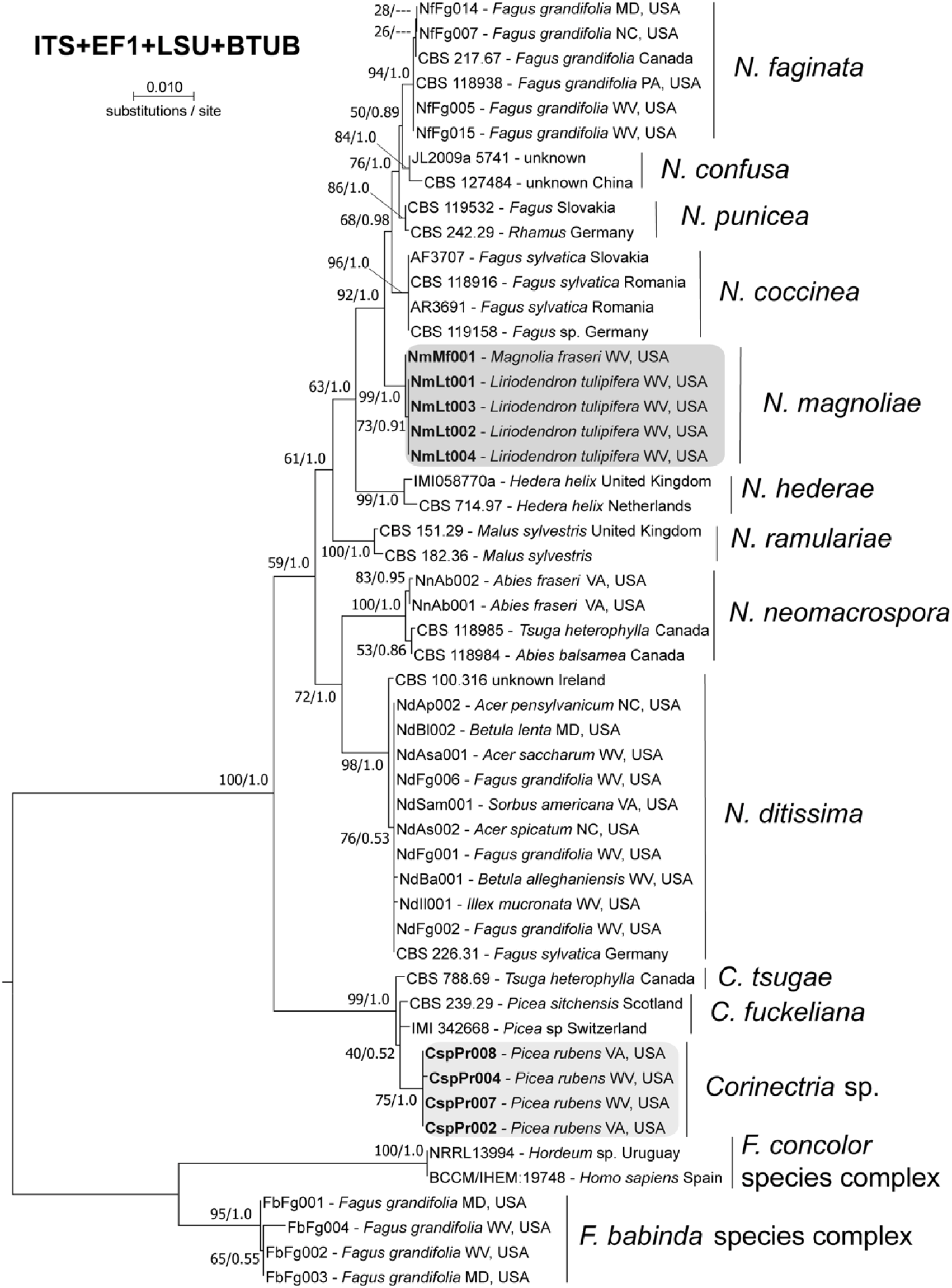
Four-gene (ITS, BTUB, ACTIN, EF1) concatenated phylogeny tree of *Neonectria* spp., *Corinectria* spp. and outgroups. Topology and branch lengths are from the ML analysis. For each node supported in the ML analysis, bootstrap support and posterior probabilities are indicated (ML/BI).

**Fig. 5:**
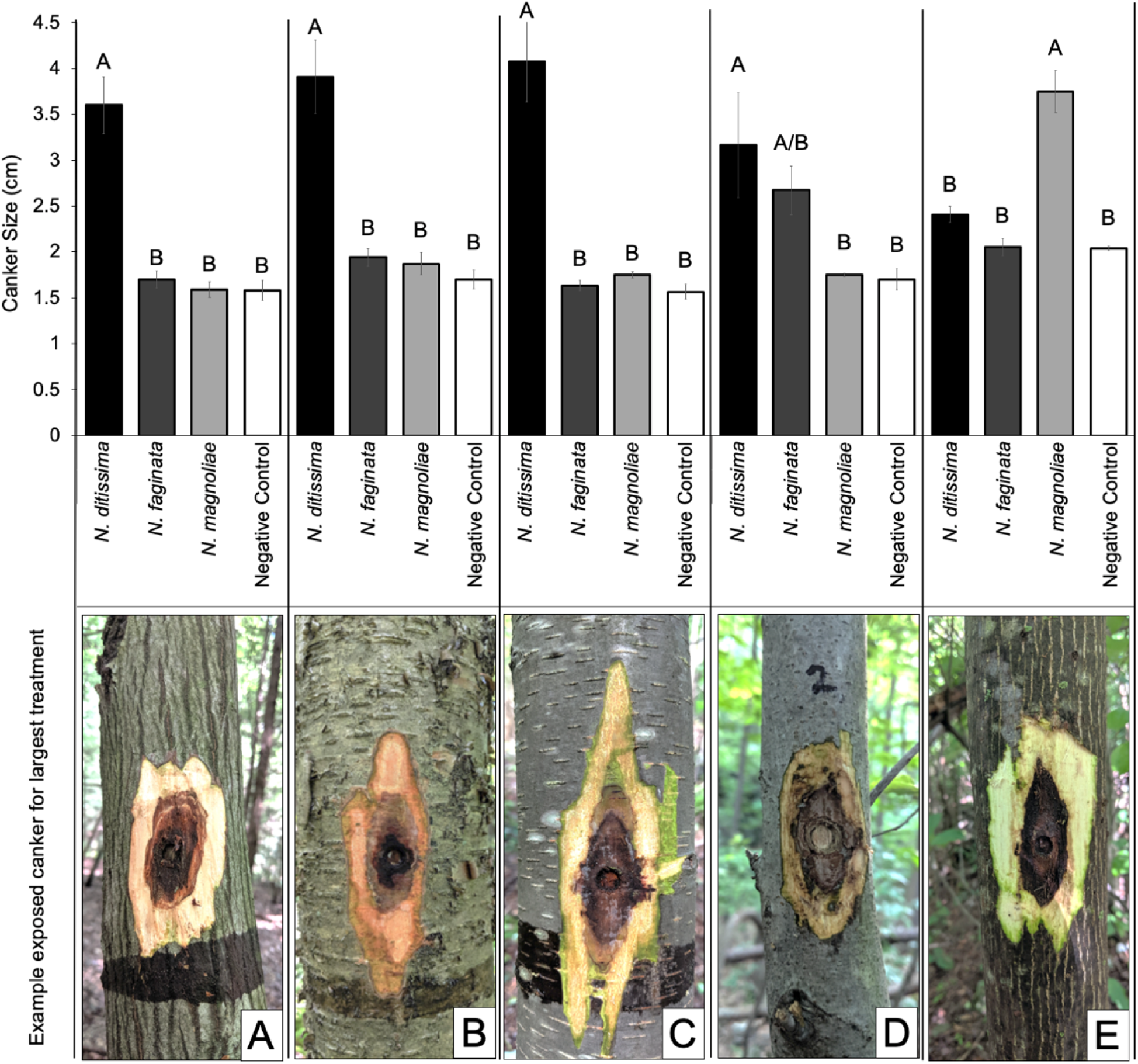
Pathogenicity results and representative photos for each host species tested: A) striped maple (*Acer pensylvanicum*), B) yellow birch (*Betula alleghaniensis*), C) black birch (*B. lenta*), D) American beech (*Fagus grandifolia*), and E) tulip poplar (*Liriodendron tulipifera*). Letters designate significant differences at p < 0.05. All sample sizes are 6 trees, except for American beech (n = 5).

**Fig. 6:**
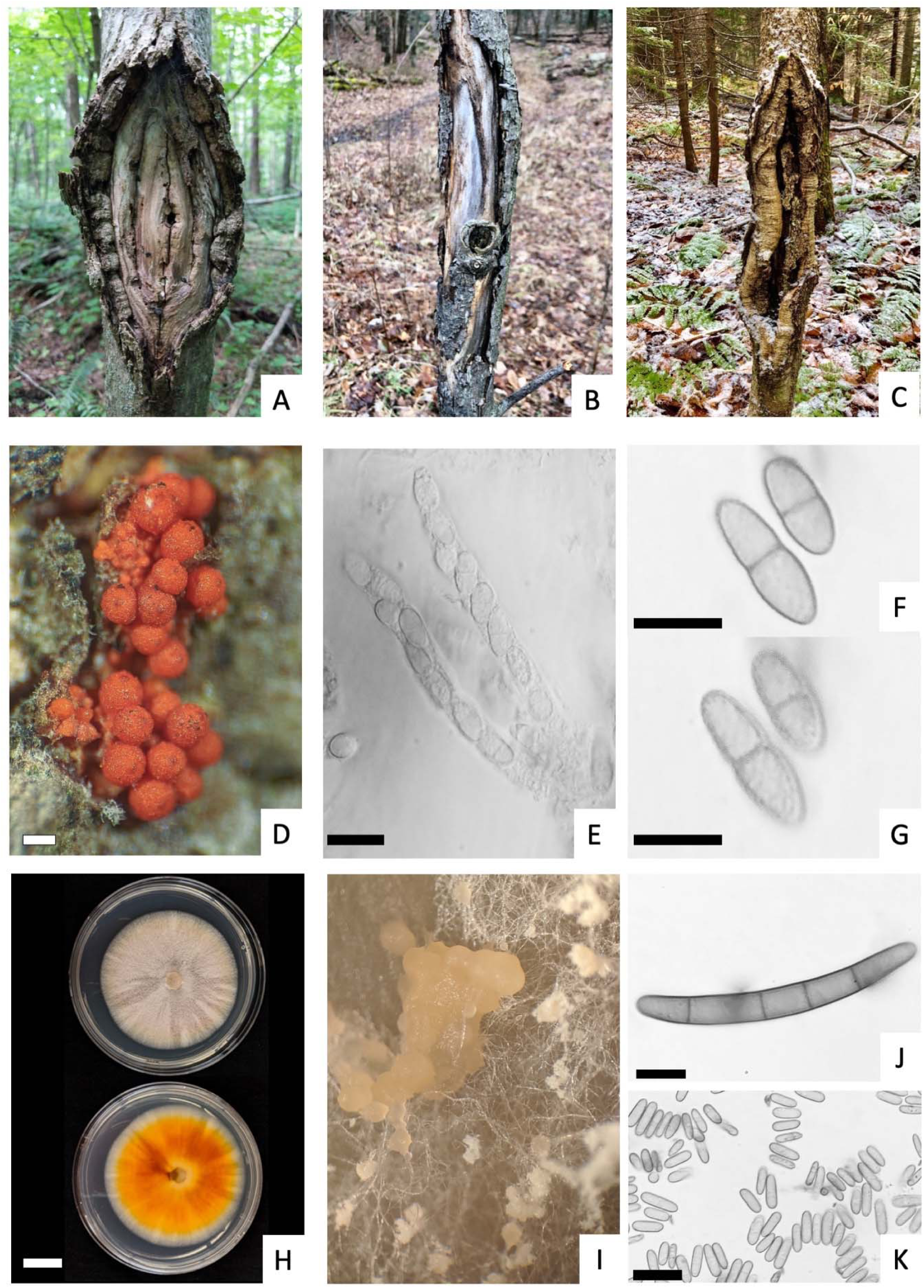
*Neonectria magnoliae*: A) Example canker on *Liriodendron tulipifera* with perennial target canker morphology; B) Example canker on *L. tulipifera* exhibiting less distinct canker morphology; C) Example canker on *Magnolia fraseri*; D) *N. magnoliae* perithecia erupting from *L. tulipfera* bark tissues; E) Asci containing eight ascospores each; F) Ascospores; G) Ornamented surface of ascospores; H) Top and bottom of 7-day old culture photos on PDA; I) Creamy microconidial sporodochia and aerial conidiophores; J) Macroconidia with four septations; K) Microconidia of varying lengths. Scale bars = 250 µm for panel D; 20 µm for panels E, K; 10 µm for panels F, G, J; 20 mm for panel H.

Three members of Nectriaceae were also confirmed on conifer hosts. These included *Thyronectria balsamea* ([Cooke & Peck] Seeler) on balsam fir (*Abies balsamea* (L.) Mill.) and *Neonectria neomacrospora* ([C. Booth & Samuels] Mantiri & Samuels) on Fraser fir (*Abies fraseri* (Pursh) Poir.) (Fig. 3E and 3C, respectively). Additionally, a novel *Corinectria* sp. on red spruce (*Picea rubens* Sarg.) was recovered in this survey across high elevation spruce forests in two states (Fig. 3D; Fig. 4).

In addition to the previously mentioned nectriaceous fungi recovered from perithecia on symptomatic beech trees and nearby co-occurring trees, a member of the *Fusarium babinda* species complex (FBSC) was commonly isolated from wood tissues of BBD-confirmed American beech trees (31.8% of bark samples), but it was lacking fruiting bodies (Supplemental Fig. 1; Supplemental Table 4). To further investigate this finding, five sites without BBD were identified, and American beech bark tissues were sampled and processed. FBSC was recovered from all confirmed BBD sites but was never recovered from non-BBD sites. Aside from *Fusarium babinda, N. faginata* and *N. ditissima* were occasionally recovered from beech bark tissues collected but only within BBD sites (Supplemental Table 4).

### 3.2. Phylogenetic Analyses

Phylogenetic analyses were performed for single genes (ITS, TEF1, TUB, LSU) and for a four-gene concatenated sequence to infer relationships among *Neonectria* and *Corinectria* species recovered in the survey. These analyses were supported with the addition of sequence data from several reference strains for both genera (Table 3). Representatives of the *Fusarium concolor* and *Fusarium babinda* species complexes were chosen to serve as outgroup taxa that could confirm the identity of *Fusarium babinda* isolates that were frequently recovered in this study from collected bark tissue samples.

The following results are summarized for relationships with >70% bootstrap support within the concatenated phylogeny (Fig. 4). *Neonectria* and *Corinectria* were resolved as strongly supported (100%/1.0) sister genera. All included *Neonectria* species were genealogically exclusive (>83%/1.0). *N. ditissima* isolates recovered from diverse host species formed a monophyletic clade with strong support (98%/1.0).*Neonectria neomacrospora* was sister to *N. ditissima* (72%/1.0) and isolates recovered in this survey were sister to selected NCBI reference sequences.

Isolates identified as *Nectria magnoliae* formed a monophyletic clade within *Neonectria* (99%/1.0) and was sister to a clade containing *N. faginata, N. coccinea, N. confusa*, and *N. punciea* (92%/1.0). The one isolate recovered from *Magnolia fraseri* (NmMf001) was divergent from all other *N. magnoliae* isolates recovered from *L. tulipifera* (73%/0.91). All single-gene trees available in Supplemental Fig. 2.

### 3.3. Pathogenicity Assay

A field pathogenicity assay was conducted to further characterize *N. magnoliae* and test the pathogenicity of *N. faginata* and *N. ditissima* isolates originating from American beech on alternative host species including those not previously included in cross pathogenicity assays. Overall, *N. ditissima* produced significantly larger cankers (p < 0.05) than all other treatments on striped maple, yellow birch, and black birch (Fig. 5A-C, respectively). *N. faginata* and *N. magnoliae* failed to produce cankers significantly larger than the negative control on any of the aforementioned hosts. On American beech, *N. ditissima* produced larger cankers than *N. magnoliae* and the negative control, but *N. ditissima* and *N. faginata* canker sizes were not significantly different (p = 0.71) (Fig. 5D). *N. faginata* produced a larger canker than *N. magnoliae*, although not statistically significant, and negative control on American beech (p = 0.19 and 0.23, respectively). *N. magnoliae* produced a significantly larger canker on tulip poplar than all other treatments (p < 0.0001) (Fig. 5E). The treatment isolate was recovered from all respective inoculations.

### 3.4. Morphological Characterization of *Neonectria magnoliae*

The morphological features of *Neonectria magnoliae* are summarized here and illustrated in Fig. 6. Cankers produced by *N. magnoliae* on tulip poplar (Fig. 6A) were reminiscent of the perennial target cankers produced by *N. ditissima*, but these cankers often appeared irregular and less descript, especially for infections on *M. fraseri* (Fig. 6B-C). Perithecia occur singly or in aggregates on bark tissue or directly on exposed wood surrounding the outer canker margins and typically emerge from thin stroma tissues as reddish-brown globous body with a distinct ostiole then fade to brown as they age (Fig. 6D). Asci appear truncated and bear eight ascospores (Fig. 6E). Ascospores ((11.3-) 13.1 – 15.3 (−17.7) µm x (4.0-) 5.6 – 7.4 (−9.1) µm) are uniseptate with rounded-ends, constricted at the septum, hyaline, and warty on the surface (Fig. 6F-G). *N. magnoliae* cultures have a white surface with an orange-red subsurface after 10 d on PDA (Fig. 6H). Microconidial sporodochia and condiophores are regularly produced in culture (Fig. 6I). Sporodochia are slimy masses with a cream to buff color. Conidiophores appear white and are penicillately branched. Microconidia ((5.7-) 8.6 – 11.0 (−13.7) µm x (3.0-) 4.0 – 5.3 (−6.4) µm) are hyaline, straight, and aseptate with rounded ends (Fig. 6 K). Microconidia are abundantly produced in culture on aerial conidiophores and within sporodochia. Macroconidia ((42.1-) 47.5 – 61.7 (−67.3) µm x (5.5-) 6.8 – 8.4 (−9.4) µm) are primarily 4-5 septate, cylindrical with rounded ends, slightly curved, and hyaline (Fig. 6J). Macroconidia, while they do occur, are found less frequently in sporodochia.

#### Taxonomy

***Neonectria magnoliae*** (Lohman & Hepting) C.M. Stauder & M.T. Kasson comb. nov.

≡ *Nectria magnoliae* M.L. Lohman & Hepting, Lloydia 6: 91. 1943.

Similar to *Neonectria coccinea*; ascospores 13.1 – 15.3 × 5.6 – 7.4 µm with regularly scattered warts; Microconidia aseptate; Macroconidia rare except for isolates from *Magnolia fraseri*. When present slightly curved 4-5 septate.

##### Holotype

UNITED STATES. West Virginia: Richwood. Civilian Conservation Corps Camp Woodbine (currently Camp Richwood) along the Cranberry River, on bark of almost dead *Liriodendron tulipifera*, Nov. 21, 1934, M. L. Lohman, BPI 552527 (No. 64184) (Fig. 7).

**Fig. 7:**
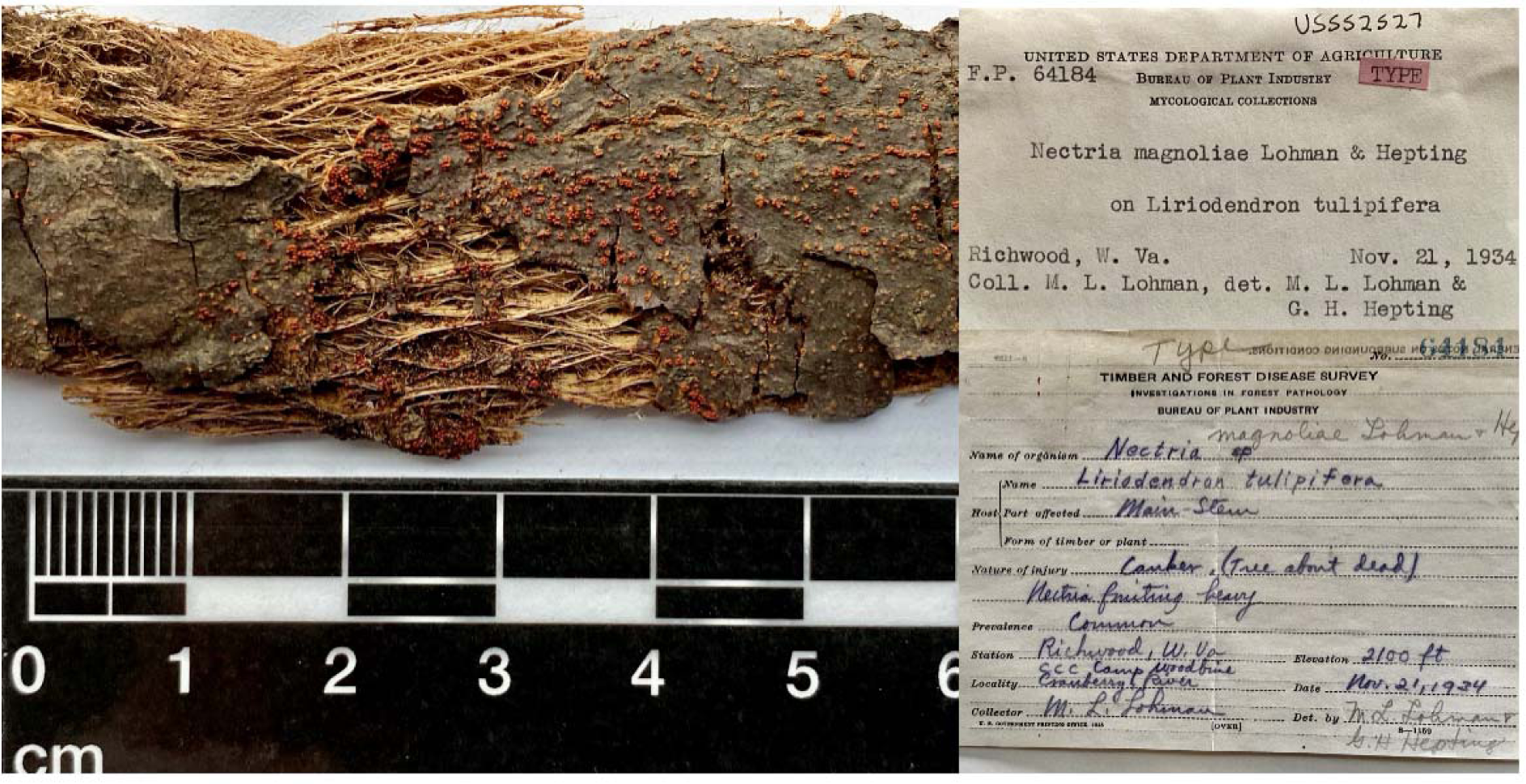
*Neonectria magnoliae* holotype (BPI 552527 [No. 64184]) submitted by Lohman and Heptig in 1934. Sample consists of *N. magnoliae* fruiting on bark of tulip poplar (*Liriodendron tulipifera*) collected in Richwood, West Virginia, USA. Photos provided by Lisa A Castlebury, Ph.D, Acting Research Leader and Collections Director, USDA ARS MNGDBL.

##### Specimens examined

UNITED STATES. North Carolina: Asheville. Bent Creek Experimental Forest, on cankered *Liriodendron tulipifera*, October 12, 1934, M. L. Lohman, BPI 552527 (No. 64187; CBS 380.50). MB#288753. ITS and RPB2 sequences available in NCBI Genbank, accessions <u>**MH856671**</u> and <u>**HQ897713**</u>, respectively. West Virginia: Monongalia County. Snake Hill Wildlife Management Area, on cankered *L. tulipifera*, October 1, 2018, C. M. Stauder and M. T. Kasson, NmLt003, NmLt004, ITS sequences available in NCBI Genbank, accessions <u>**TBD**</u>, <u>**TBD**</u>, respectively. Tucker County. Fernow Experimental Forest, on cankered *L. tulipifera*, July 12, 2018, C.M. Stauder, A. M. Macias, M. T. Kasson, NmLt001, NmLt002, ITS sequences available in NCBI Genbank, accessions <u>**TBD**</u>, <u>**TBD**</u>, respectively. Randolph County. Gaudineers Knob, on cankered *Magnolia fraseri*, M.T. Kasson, NmMf001, ITS sequences available in NCBI Genbank, accessions <u>**TBD**</u>.

##### Habitat and distribution

on bark and exposed wood around cankers on living and dead *Liriodendron tulipifera* L. in Connecticut, Ohio, West Virginia, Virginia, Tennessee and North Carolina, *Magnolia fraseri* Walt. trees in West Virginia and Tennessee, and *M. tripetala* L. in West Virginia. Trees include both overtopped and overstory trees. Often found in forested stands co-occurring with other hosts harboring conspicuous *N. ditissima* target cankers.

##### Descriptions and illustrations

See Lohman and Watson (1943).

## 4. Discussion

In this study, we sampled and characterized the diversity of Nectriaceae across the central Appalachian Mountains. Over the course of the survey, ten species of Nectriaceae belonging to *Bionectria, Corinectria, Cosmospora, Fusarium, Neonectria, Thelonectria*, and *Thyronectria* were recovered from twelve tree hosts spanning 17 sites across six states (Table 1; Fig. 3). The limited recovery of *Cosmospora obscura, Neonectria neomacrospora, Thelonectria balsamea*, and *Thyronectria veuillotiana* limits our ability to explore their ecology but follow-up surveys/studies on these species could provide further insight into lifestyles of these fungi.

Other recovered members of Nectriaceae were either widespread or found in sufficient numbers at fewer locations, thereby providing adequate resolution and/or insight into their ecology. These species included the following: *Neonectria magnoliae* from symptomatic tulip poplar and Fraser magnolia; *N. ditissima* from eight hosts; *N. faginata, Bionectria ochroleuca*, and *Fusarium babinda* from *American beech*; and a novel *Corinectria* sp. from red spruce (Table 1; Fig. 3).

*Neonectria magnoliae* was first described as *Nectria magnoliae* by Lohman and Watson (1943) as having some overlapping characteristics of both *Neonectria ditissima* (formerly *Nectria galligena*) and *Neonectria coccinea* (formerly *Nectria coccinea*), but also with notable morphological features including progressive changes in coloration of perithecia and notable spore size differences. Many former members of the genus Nectria have since been reclassified into different genera (Brayford et al., 2004; Castlebury et al., 2006; Mantiri et al., 2001; Rossman et al., 1999). However, none of these studies considered *Nectria magnoliae*, and therefore, its taxonomy was never properly resolved with one erroneous exception: Castlebury and colleagues (2006) included one isolate of *Nectria magnoliae* from tulip poplar from Tennessee in a phylogenetic study of *Neonectria*. These authors concluded that *N. magnoliae* was a synonym of *N. ditissima*, based on the data available at the time. However, based on BLASTn searches against NCBI Genbank and pairwise blasting of our *N. magnoliae* isolates with *N. ditissima* strain CBS 118919, it’s clear that both *N. magnoliae* and *N. ditissima* infect tulip poplar and strain CBS 118919, sequenced by Castebury et al. (2006) was in fact *N. ditissima*. NCBI BLASTn searches with ITS and RPB2 sequences generated for our field-collected *N. magnoliae* isolates had 98.4%-100% sequence similarity with 77-100% sequence coverage with a paratype of *Nectria magnoliae* (CBS 380.50 Genbank Accessions NR_160076 and HQ897713, respectively) (Supplemental Table 3). Additionally, the four-gene concatenated phylogeny placed *N. magnoliae* isolates as a monophyletic ingroup within *Neonectria* (Fig. 4).

Ascospore measurements and general morphological descriptions provided in the type description for *N. magnoliae* (Lohman and Watson, 1943) were comparable with those observed in this study (Fig. 6). Notable was the infrequency of macroconidia from fresh cultures on general growth media. Interestingly, this trend did not hold for all isolates as several of the *N. magnoliae* from Fraser magnolia produced abundant macroconidia even in week-old cultures. The only isolate of *N. magnoliae* from magnolia also showed some sequence divergence from all tulip poplar isolates (Fig. 4). Further characterization of magnolia and tulip-polar isolates are needed to confirm whether these notable differences are biologically significant. The included pathogenicity trial confirmed the pathogenicity of *N. magnoliae* on tulip poplar but not on other hosts tested (Fig. 5). Interestingly, *N. ditissima* did not produce cankers significantly different from the negative control on tulip-poplar despite the previous work by Castlebury et al. (2006) confirming tulip-poplar as a host of *N. ditissima* (Fig. 5). Earlier work by Lohman and Watson (1934) also detail cross-pathogenicity experiments in North Carolina where diverse *N. ditissima* strains from various hosts caused significantly larger cankers on tulip poplar compared to *N. magnoliae*. Follow-up studies are needed for cross pathogenicity tests of N. *magnoliae* from different hosts as well as *N. ditissima* isolates from tulip poplar. Together, these results provide evidence to support the transfer of *Nectria magnoliae* into the genus *Neonectria*.

*N. ditissima* and *N. faginata* represented a majority of isolates recovered during this study (Table 1). Although this study did not uncover cryptic native plant reservoirs of *N. faginata, N. ditissima* was confirmed from eight plant hosts including three of which may represent the first reports: *Acer spicatum, Ilex mucronata*, and *Sorbus americana* (Table 1). Interestingly, not all hosts had characteristic perennial target cankers (Fig. 2). Instead, *Acer pensylvanicum* and *Ilex mucronata* exhibited vascular cambium necrosis and perithecia production without an apparent outward host response (*e.g*. callous ridges, bark malformation). Perithecia production also varied across hosts with black birch cankers consistently yielded some perithecia and mountain ash cankers rarely yielding perithecia.

Cross-pathogenicity assays among *N. ditissima* strains demonstrated the lack of host specificity between isolates recovered from American beech or alternate host species (Fig. 5). Additionally, a recent study confirmed the lack of mating barriers among *N. ditissima* strains recovered from varying host tree species (Stauder et al., 2020). Together, these results provide additional observational evidence describing the extent of the generalist nature of *N. ditissima* as a ubiquitous phytopathogen in Eastern North America capable of infecting many hardwood species.

*Neonectria faginata* was the most recovered member of the Nectriaceae in this study, though it was exclusive to American beech (Table 1). The identification of a cryptic reservoir may have provided additional evidence supporting the nativity of *N. faginata* to Eastern North America. Alternatively, *N. faginata* may be an exotic species simply capable of infecting as-yet-unidentified host species given this remains a possibility for American beech. Either way, no evidence of an alternate host for *N. faginata* was uncovered in this survey. Follow-up studies should focus on cankers lacking perithecia and or asymptomatic host tissue similar to what has been previously described for epiphytic *N. coccinea* on European beech (Chapela and Boddy, 1988; Hendry et al., 2002).

Previous assertions of the eventual dominance of *N. faginata* in the BBD pathosystem were confirmed across our study sites (Houston, 1994; Kasson and Livingston, 2009). In total, *N. faginata* was recovered from 93.7% of BBD samples collected in this survey (from 13 of 13 BBD sites) while *N. ditissima* was only recovered from 4.2% of BBD samples (from two of the 13 BBD sites) (Table 1).

Nevertheless, *N. ditissima* still appears to play a minor role in BBD in its more advanced stages. The pathogenicity trial demonstrated the potentially increased pathogenicity of *N. ditissima* on American beech when compared to *N. faginata* (Fig. 5). While the interaction of the scale insect may have significantly altered the host’s physiology and thus, defense responses to fungal infections, these results appear to indicate that an increased virulence of *N. faginata* may not be responsible for its dominance in the BBD pathosystem.

One additional possibility may be related to its production of perithecia and the resulting inoculum potential. For example, taxonomic descriptions of *N. ditissima* and *N. faginata* illustrate differences in their production of perithecia. In general, *N. ditissima* is described as bearing fewer, scattered perithecia while *N*. faginata can be found to produce larger aggregates in higher densities (Castlebury et al., 2006; Lohman and Watson, 1943). Sampling of *N. ditissima* perithecia across eight hosts generally supports these previous observations (Table 1). Seasonal differences in fruiting could explain differential abundances in these two fungi, but *N. ditissima* isolates were recovered from beech and non-beech hosts during the same sampling periods in which sampling of BBD trees resulted in high abundances of *N. faginata*. Therefore, seasonality of fruiting does not appear to have been an apparent factor in this study. Further, several studies report ascospores as the predominant spore type in the environment and more specifically, the dominant infective spore type (Crane et al., 2009; Lortie and Kuntz, 1963). While these are general descriptive observations and not quantitative comparisons, differences in fruiting habits may explain the eventual abundance of *N. faginata* due to a significantly increased inoculum potential.

*Bionectria ochroleuca* was also recovered from BBD impacted beech trees at two total sites in West Virginia and North Carolina, representing 2.1% of all perithecia sampled from beech (Table 1; Fig. 3). This fungus had been previously implicated in BBD although its geographic range was not well studied (Houston et al., 1987). Here we update the known distribution of this fungus to include West Virginia and North Carolina. Our observations of this fungus on successfully colonized trees supports previous studies, which indicate this fungus is mycoparasitic (Barnett and Lilly, 1962; Jager et al. 1979; Turhan, 1993). Isolates of this fungus were also recovered from American basswood in North Carolina during our surveys (Table 1). This appears to be the first report of this fungus on American basswood in the U.S.

An additional fungus recovered consistently from beech with BBD was a member of the *Fusarium babinda* species complex (FBSC), although it did not produce perithecia, like all the other Nectriaceae recovered in this study. Instead, members of the FBSC often found colonizing the base of *Neonectria* perithecia or from necrotic tissue around *Neonectria* fruiting bodies. After its discovery in single spore plates from *Neonectria* perithecia, follow-up studies using micro-sampled bark tissue revealed this fungus was present in 31.8% of BBD-positive bark samples at 100% of BBD sampled sites but not present in any beech bark samples taken from healthy asymptomatic trees outside confirmed BBD epicenters (Supplemental Table 4). Its prevalence from BBD impacted stands, but lack of previous reporting, indicates either a localized (albeit unclear) role or novel component in BBD pathosystem. However, a re-assessment of raw data and culture photos taken by the senior author during 2005 – 2006 as part of his BBD M.S. research in Maine (Kasson and Livingston, 2009) uncovered a *Fusarium* with identical morphology to *F. babinda* in three of three sites sampled for which culture data was collected (Supplemental Fig. 1; See Kasson and Livingston, 2007, Fig. 17A). This clearly indicates a more widespread association with BBD and one which has not been observed previously despite over 130 years of research on this important pathosystem. One possibility may be that *F. babinda* is insecticolous, interacting with the scale as populations decline and as *Neonectria* spp. colonize stems, which is likely given this fungus has been recovered from both gypsy moth and hemlock woolly adelgid populations in the eastern United States (Jacobs-Venter et al., 2018). Another plausible, although not mutually exclusive hypothesis based on the observations of sporulation at the base of viable and nonviable perithecia, is that *F. babinda* is a facultative hyperparasite of *Neonectria*. Clearly, more work is needed to test phytopathogenicity and entomopathogenicity, and determine its dominant and facultative lifestyles. Preliminary testing of its ability to degrade tannic acid, a proxy assay for lignin degradation (Kasson et al., 2016) clearly shows that it has strong lignin and cellulose degrading activity.

Finally, in our repeated attempts to uncover non-conventional hosts of *N. faginata*, we discovered a putatively novel *Corinectria* sp. on red spruce in high elevation spruce-fir forests in close proximity to BBD epicenters in VA and WV (Table 1; Fig. 3). Interestingly, the fungus appears to be associated with dead trees but the results of those studies and in-depth investigations into the biology and ecology of this putatively novel species are forthcoming. However, the *Corinectria* sp. recovered formed a single clade and was divergent from *C. fuckeliana* and *C. tsugae* in the concatenated phylogeny (Fig. 4). The third described *Corinectria* species, *C. constricta*, was not included in the concatenated phylogeny due to limited availability of sequence data, but based on EF1, isolates recovered in this study were divergent from *C. constricta* as well. While this appears to be evidence of a novel species of *Corinectria*, further investigation is needed.

Taken together, the results of this broad study emphasize the sheer diversity of Nectriaceae present in the central Appalachian Mountains. Still, this is undoubtedly a gross underestimation of known and novel fungi as this survey intentionally targeted members of the Nectriaceae with viable and conspicuous perithecia. Homothallic Nectriaceae as well as exclusively anamorphic fungi were largely excluded with the exception of *F. babinda*. The insights gained here into other fungi of the BBD pathosystem opens many opportunities for further investigation. Specifically, what is the dominant lifestyle of *B. ochroleuca*, and what role do members of the FBSC actually play in BBD where it appears to dominate tissues recently colonized by scale insects and/or *Neonectria* fungi? These studies provide an important baseline for how fungal communities assemble, change over time, and affect the forests of the Appalachian Mountains.

## 5. Acknowledgements

Special thanks to Braley Burke and Amy Metheny for their assistance with single spore isolations and Wyatt, Oilver, and Andrew Kasson for their help conducting filed collections. Thank you to Dr. Lisa Castlebury, USAD ARS Mycology & Nematology Genetic Diversity & Biology Lab Collections Director, for providing photos of the type specimen of *N. magnoliae*. C.M.S. was supported, in part, by the WVU Outstanding Merit Fellowships for Continuing Doctoral Students, N.M.U. by WVU SURE program supported by the NSF-funded KY-WV LSAMP Alliance, and M.T.K. by funds from West Virginia Agricultural and Forestry Experiment Station, Morgantown, WV.

