## Supplemental Figures for "Resolving host and species boundaries for perithecia-producing nectriaceous fungi across the central Appalachian Mountains"

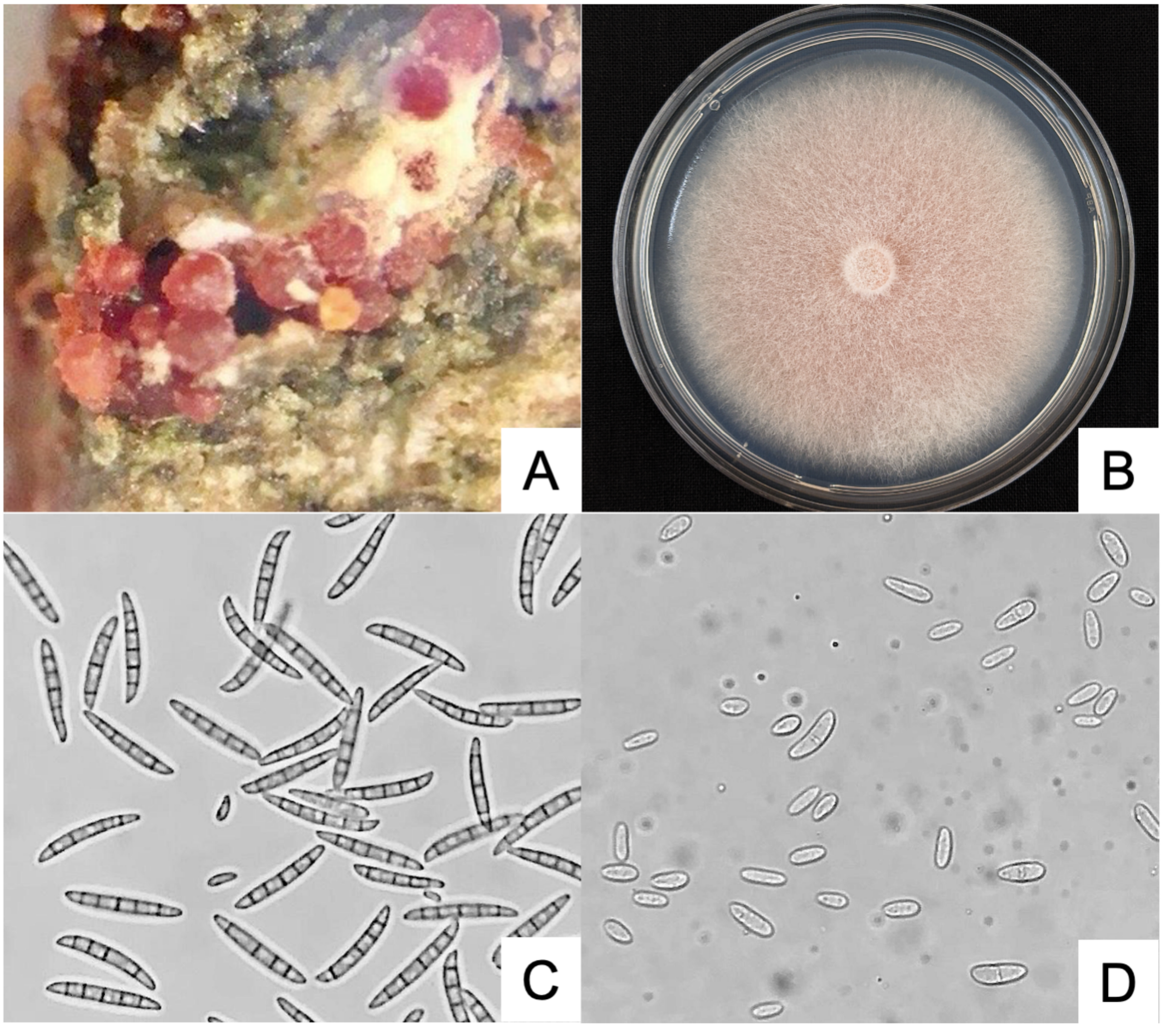


**Supplemental Fig. 1:** Morphological characteristics of *Fusarium babinda*: A) *F. babinda* sporulation on *Neonectria* sp. perithecia; B) 7-day old culture on PDA; B) sporodochia bearing macro- and micro-conidia; C) macroconidia; D) microconidia

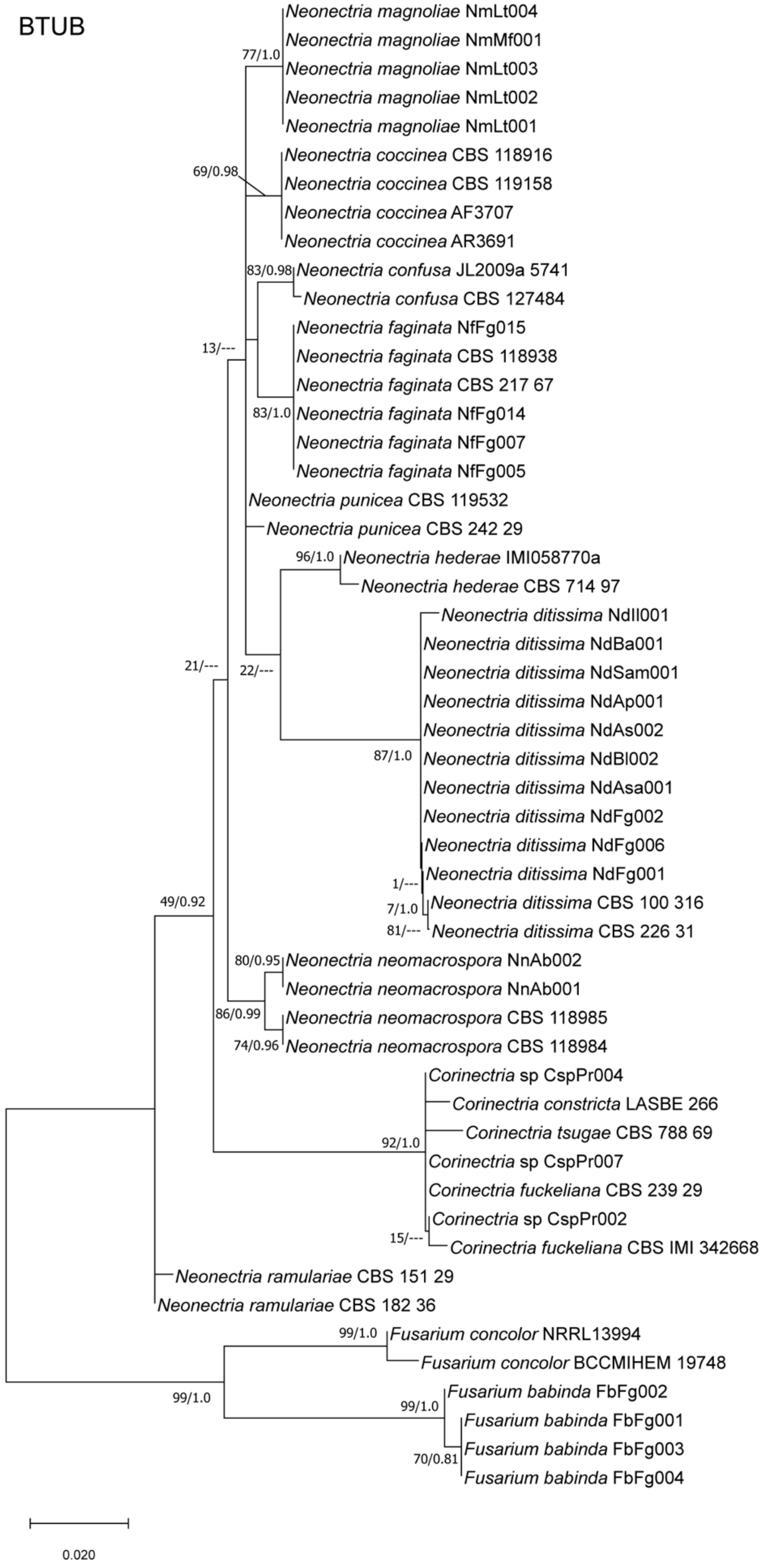

BTUB
Neonectria magnoliae NmLt004
Neonectria magnoliae NmMf001
Neonectria magnoliae NmLt003
Neonectria magnoliae NmLt002
Neonectria magnoliae NmLt001
Neonectria coccinea CBS 118916
Neonectria coccinea CBS 119158
Neonectria coccinea AF3707
Neonectria coccinea AR3691
Neonectria confusa JL2009a 5741
Neonectria confusa CBS 127484
Neonectria faginata NfFg015
Neonectria faginata CBS 118938
Neonectria faginata CBS 217 67
Neonectria faginata NfFg014
Neonectria faginata NfFg007
Neonectria faginata NfFg005
Neonectria punicea CBS 119532
Neonectria punicea CBS 242 29
Neonectria hederae IMI058770a
Neonectria hederae CBS 714 97
Neonectria ditissima NdIl001
Neonectria ditissima NdBa001
Neonectria ditissima NdSam001
Neonectria ditissima NdAp001
Neonectria ditissima NdAs002
Neonectria ditissima NdBl002
Neonectria ditissima NdAsa001
Neonectria ditissima NdFg002
Neonectria ditissima NdFg006
Neonectria ditissima NdFg001
Neonectria ditissima CBS 100 316
Neonectria ditissima CBS 226 31
Neonectria neomacrospora NnAb002
Neonectria neomacrospora NnAb001
Neonectria neomacrospora CBS 118985
Neonectria neomacrospora CBS 118984
Corinectria sp CspPr004
Corinectria constricta LASBE 266
Corinectria tsugae CBS 788 69
Corinectria sp CspPr007
Corinectria fuckeliana CBS 239 29
Corinectria sp CspPr002
Corinectria fuckeliana CBS IMI 342668
Neonectria ramulariae CBS 151 29
Neonectria ramulariae CBS 182 36
Fusarium concolor NRRL13994
Fusarium concolor BCCMIHEM 19748
Fusarium babinda FbFg002
Fusarium babinda FbFg001
Fusarium babinda FbFg003
Fusarium babinda FbFg004
77/1.0
69/0.98
83/0.98
13/---
83/1.0
96/1.0
21/---
22/---
87/1.0
1/---
7/1.0
81/---
49/0.92
80/0.95
86/0.99
74/0.96
92/1.0
15/---
99/1.0
99/1.0
99/1.0
70/0.81
0.020

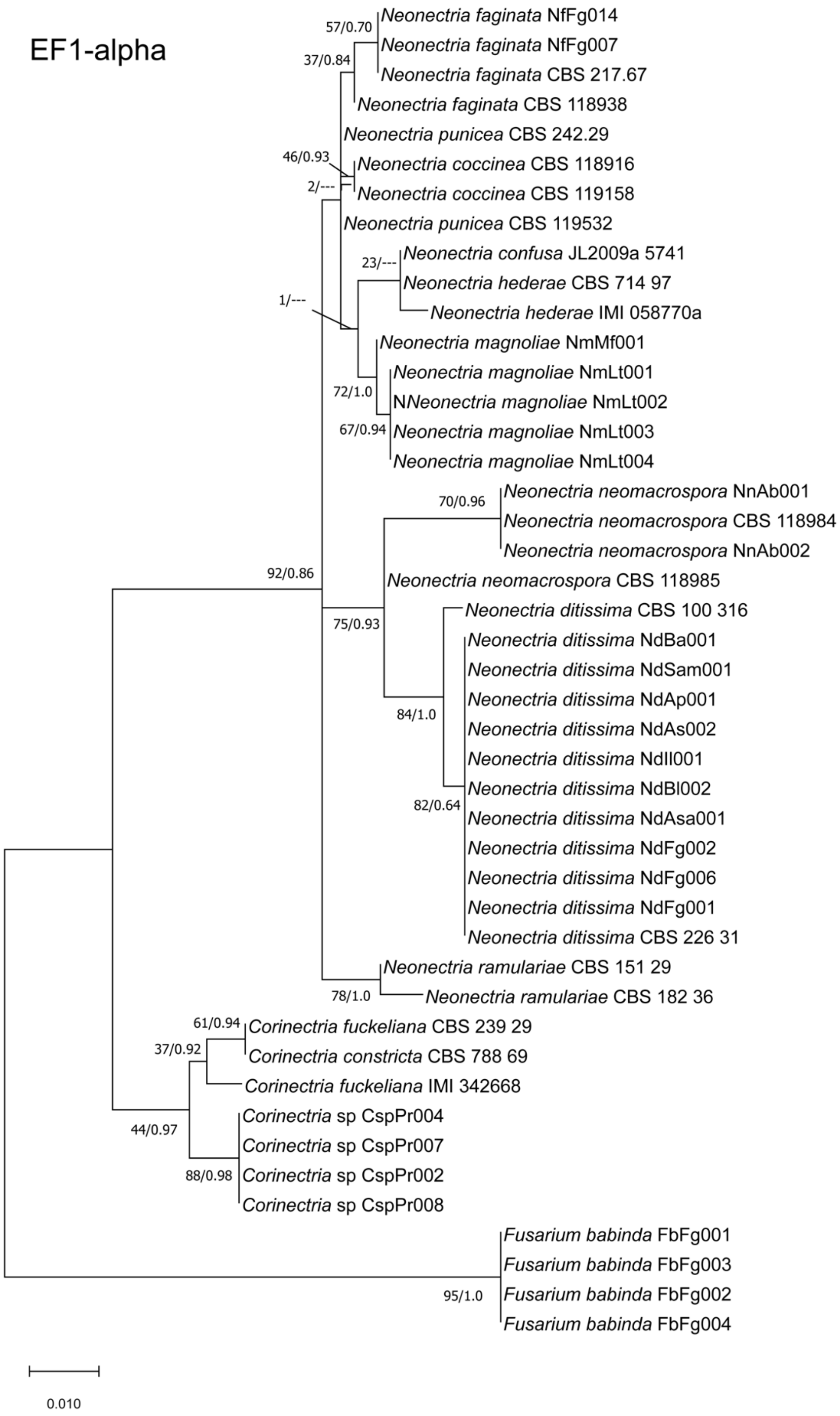
EF1-alpha
Neonectria faginata NfFg014
Neonectria faginata NfFg007
Neonectria faginata CBS 217.67
Neonectria faginata CBS 118938
Neonectria punicea CBS 242.29
Neonectria coccinea CBS 118916
Neonectria coccinea CBS 119158
Neonectria punicea CBS 119532
Neonectria confusa JL2009a 5741
Neonectria hederae CBS 714 97
Neonectria hederae IMI 058770a
Neonectria magnoliae NmMf001
Neonectria magnoliae NmLt001
NNeonectria magnoliae NmLt002
Neonectria magnoliae NmLt003
Neonectria magnoliae NmLt004
Neonectria neomacrospora NnAb001
Neonectria neomacrospora CBS 118984
Neonectria neomacrospora NnAb002
Neonectria neomacrospora CBS 118985
Neonectria ditissima CBS 100 316
Neonectria ditissima NdBa001
Neonectria ditissima NdSam001
Neonectria ditissima NdAp001
Neonectria ditissima NdAs002
Neonectria ditissima NdIl001
Neonectria ditissima NdBl002
Neonectria ditissima NdAsa001
Neonectria ditissima NdFg002
Neonectria ditissima NdFg006
Neonectria ditissima NdFg001
Neonectria ditissima CBS 226 31
Neonectria ramulariae CBS 151 29
Neonectria ramulariae CBS 182 36
Corinectria fuckeliana CBS 239 29
Corinectria constricta CBS 788 69
Corinectria fuckeliana IMI 342668
Corinectria sp CspPr004
Corinectria sp CspPr007
Corinectria sp CspPr002
Corinectria sp CspPr008
Fusarium babinda FbFg001
Fusarium babinda FbFg003
Fusarium babinda FbFg002
Fusarium babinda FbFg004
57/0.70
37/0.84
46/0.93
2/---
23/---
1/---
72/1.0
67/0.94
70/0.96
92/0.86
75/0.93
84/1.0
82/0.64
78/1.0
61/0.94
37/0.92
44/0.97
88/0.98
95/1.0
0.010

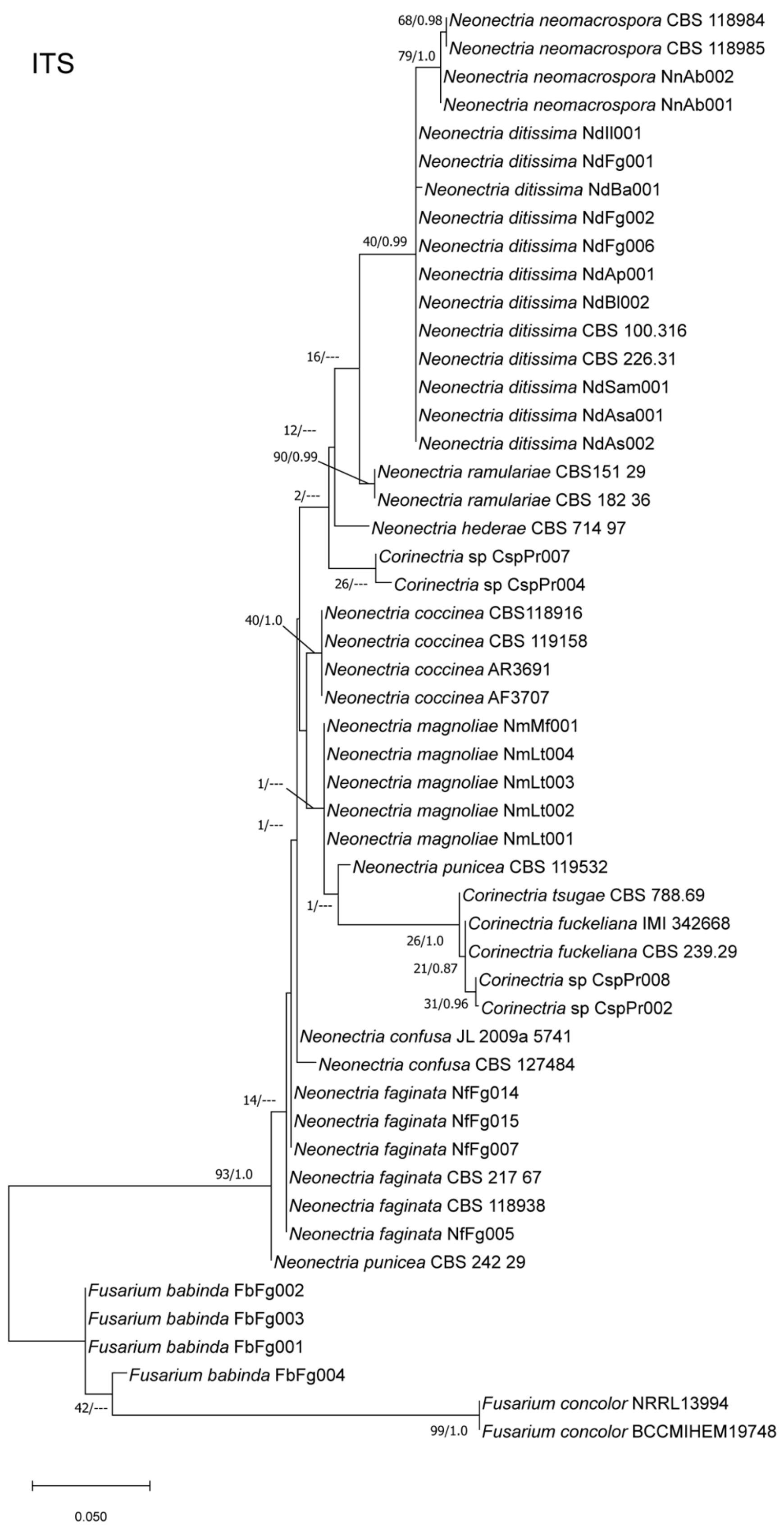
ITS
68/0.98
Neonectria neomacrospora CBS 118984
79/1.0
Neonectria neomacrospora CBS 118985
Neonectria neomacrospora NnAb002
Neonectria neomacrospora NnAb001
Neonectria ditissima NdIl001
Neonectria ditissima NdFg001
Neonectria ditissima NdBa001
Neonectria ditissima NdFg002
40/0.99
Neonectria ditissima NdFg006
Neonectria ditissima NdAp001
Neonectria ditissima NdBl002
Neonectria ditissima CBS 100.316
16/---
Neonectria ditissima CBS 226.31
Neonectria ditissima NdSam001
Neonectria ditissima NdAsa001
12/---
Neonectria ditissima NdAs002
90/0.99
Neonectria ramulariae CBS151 29
2/---
Neonectria ramulariae CBS 182 36
Neonectria hederae CBS 714 97
Corinectria sp CspPr007
26/---
Corinectria sp CspPr004
Neonectria coccinea CBS118916
40/1.0
Neonectria coccinea CBS 119158
Neonectria coccinea AR3691
Neonectria coccinea AF3707
Neonectria magnoliae NmMf001
Neonectria magnoliae NmLt004
1/---
Neonectria magnoliae NmLt003
Neonectria magnoliae NmLt002
1/---
Neonectria magnoliae NmLt001
Neonectria punicea CBS 119532
1/---
Corinectria tsugae CBS 788.69
Corinectria fuckeliana IMI 342668
26/1.0
Corinectria fuckeliana CBS 239.29
21/0.87
Corinectria sp CspPr008
31/0.96
Corinectria sp CspPr002
Neonectria confusa JL 2009a 5741
Neonectria confusa CBS 127484
14/---
Neonectria faginata NfFg014
Neonectria faginata NfFg015
Neonectria faginata NfFg007
93/1.0
Neonectria faginata CBS 217 67
Neonectria faginata CBS 118938
Neonectria faginata NfFg005
Neonectria punicea CBS 242 29
Fusarium babinda FbFg002
Fusarium babinda FbFg003
Fusarium babinda FbFg001
Fusarium babinda FbFg004
42/---
Fusarium concolor NRRL13994
99/1.0
Fusarium concolor BCCMIHEM19748
0.050

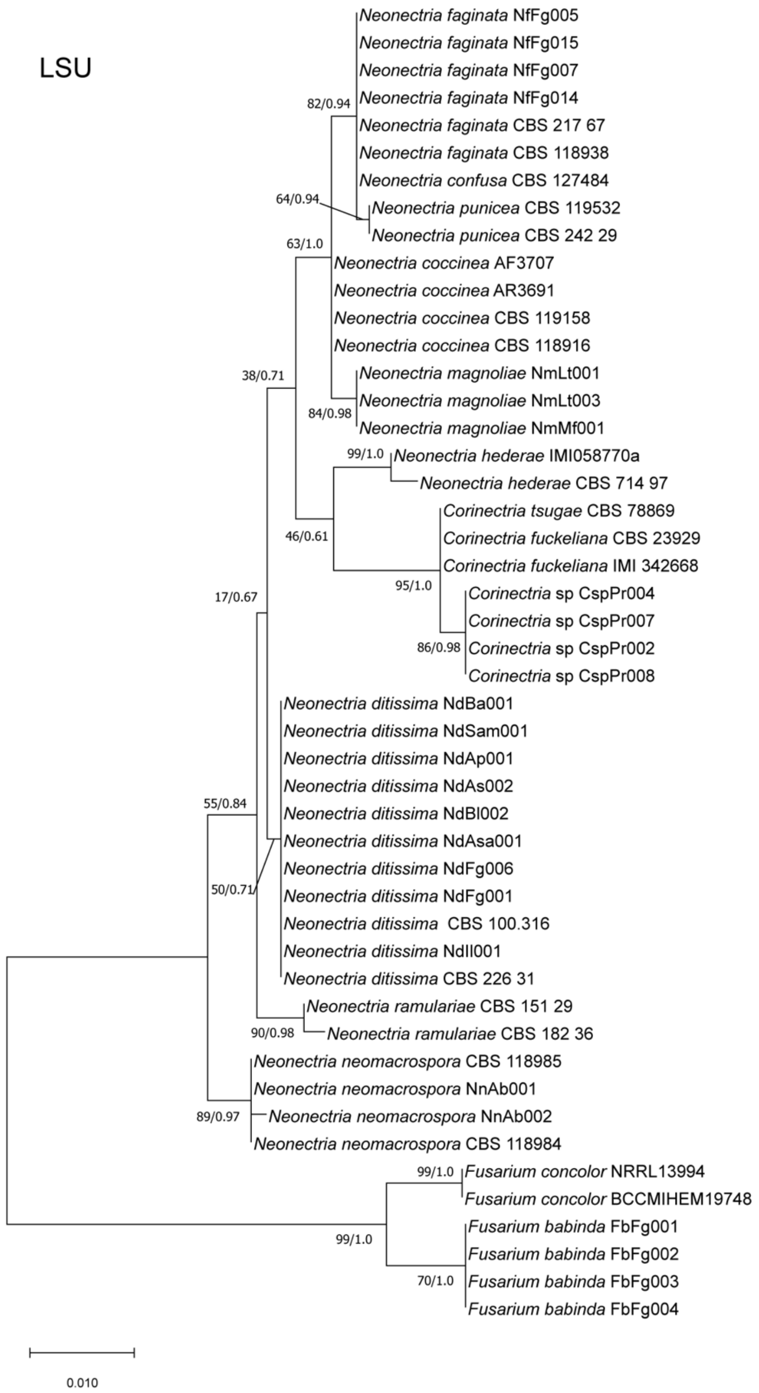


**Supplemental Figs. 2 - 5:** Single-gene (BTUB, EF1, ITS, LSU) phylogenetic tree of *Neonectria* spp., *Corinectria* spp. and outgroups. Topology and branch lengths are from the ML analysis. For each node supported in the ML analysis, bootstrap support and posterior probabilities are indicated (ML/BI).
